# Application of Elevated Atmospheric Pressure and Hypoxia Enhance Pluripotency and Stem Cell Differentiation

**DOI:** 10.1101/2024.01.19.576329

**Authors:** Zachary Pappalardo, Bryan Downie, Bruce A. Adams, James Lim

## Abstract

Physical forces regulate stem cell differentiation in-vivo, however few simple and precise methods exist to better understand this biology in-vitro. Here we describe the use of a novel bioreactor that enables addition of physical force in the form of elevated atmospheric pressure during reprogramming of human fibroblasts and culture of human induced pluripotent stem cell (iPSC) and neural stem cell (NSC) lines. We demonstrate that elevated atmospheric pressure and hypoxia can positively regulate reprogramming of human fibroblasts to iPSCs across multiple donors. Prolonged culture of iPSCs in elevated atmospheric pressure (+ 2 PSI) and 15% oxygen exhibited progressive differentiation with concomitant metabolic and epigenetic gene expression changes. Furthermore, elevated atmospheric pressure positively regulates differentiation of iPSCs to neural-ectodermal and hematopoietic lineages when combined with appropriate soluble factors and oxygen concentration. In summary, these results demonstrate the significance of applied atmospheric pressure for stem cell applications and warrants further investigation.

## Introduction

An increasing number of studies are revealing a role for micro-environmental control as it relates to stem cell maintenance and differentiation by factors such as oxygen concentration and physical force. The influence of oxygen concentration on stem cell maintenance (Yoshida et al, 2009; Mathieu et al, 2013) and differentiation (Zhu et al, 2005; Giese et al, 2010) is well-documented, and the implementation of physical force during in-vitro stem cell culture is gaining traction in the field. Methods of physical force application in stem cell culture attempt to recapitulate the stem cell niche in-vivo, including mechanical force due to three-dimensional cell-to-cell interactions, extracellular matrix (ECM) composition and stiffness, and fluid shear stress generated by the vascular system (Eiraku and Sasai, 2012; Zhong et al, 2014; Yin et al, 2016; Engler et al, 2006; Saha et al, 2008; Leipzig et al, 2009; Shimizu et al, 2008; Correia et al, 2014). Protocols for stem cell differentiation occasionally implement these forms of physical force; however, they often include only a single element of force at a time and additionally, variable inter-laboratory outcomes exist due to a lack of standardized delivery methods (Battista et al 2005; Hong et al, 2010; McKee and Chaudhry, 2017; Tworkoski et al, 2018). To address these limitations and to provide insight into a largely under-appreciated environmental variable in cell culture, we employed a novel bioreactor that can apply precise levels of physical force in the form of atmospheric pressure and oxygen that can be leveraged with available stem cell differentiation protocols. In the current study, we provide a proof-of-concept demonstrating that elevated atmospheric pressure and hypoxia can enhance somatic cell reprogramming and stem cell differentiation workflows.

Common cell culture methodology relies on incubation in static pressure and ambient oxygen concentration, which in the context of the human body are considered non-physiologic. For example, a wide range of oxygen concentration encompasses the adult human body and developing tissues during adult stem cell differentiation and embryogenesis, respectively, yet methods to recapitulate in-vivo oxygen concentration are not commonly employed in stem cell culture. What is deemed “hypoxic” in the laboratory cell culture incubator (i.e. 1-5% oxygen) more closely mimics tissue normoxia than conventional CO_2_ incubators that maintain a constant ∼18.6% oxygen concentration (Place et al, 2017). For example, physiologic oxygen concentration in the developing mammalian brain and cortical tissue is between 1 to 5% (Studer et al, 2000; Sakadzic et al, 2010). In vitro studies using hypoxic culture reveal that neurogenesis is promoted in fetal human neurons maintained in 3% O_2_ and hypoxia inducible factor (HIF) proteins can promote adult stem cell differentiation to astrocytes (Ortega et al, 2016; Yasui et al, 2017). Similarly, hematopoietic stem/progenitor cells (HSPCs) in the bone marrow niche experience oxygen concentration in the range of 1-3%, and when leveraged in vitro exhibit greater self-renewal capacity (Spencer et al, 2014; Ivanovic et al, 2004). These observations support the rationale for applying physiologic oxygen concentration during culture.

The implementation of atmospheric pressure as a physical force during cell culture has been evaluated across a wide range of systems, including batch microbial culture (Lopes et al, 2014), simulation of the vascular endothelium tissue micro-environment (Ohashi et al, 2007), disease-modeling for glaucoma and hypertension (Luo et al, 2014; Stanley et al, 2005), and induction of cancer metastasis by simulating increased interstitial fluid pressure of solid tumors (Kao et al, 2017). Like these studies, our bioreactor technology imparts physical force on cells through increased hydrostatic pressure, but by means of gas compression above the liquid layer. To date, few if any studies demonstrate the role of atmospheric pressure on pluripotent stem cell generation and differentiation, however its positive influence on adult stem cell differentiation has been reported. For example, chondrogenic and osteogenic differentiation of mesenchymal stem cells is shown to be positively regulated by elevated atmospheric pressure (Labedz-Maslowska, 2021; Zhou et al, 2014; Wagner et al, 2008) and there is a recognized need for the ability to produce functional HSPCs ex vivo for clinical use and pressure could be one of factors to achieve this (Li et al, 2021). Translating these regulatory effects of atmospheric pressure to iPSCs, which have virtually limitless differentiation potential, could enable more efficient generation of lineage-specific cell types.

To evaluate the role of atmospheric pressure during stem cell culture, we first examined its influence on reprogramming of somatic cells to iPSCs. When combined with hypoxia (1% O_2_), transient culture of fibroblasts transfected with reprogramming factors in elevated pressure (+ 5 PSI, relative to ambient) resulted in a significant increase in reprogramming efficiency across 10 unique human donors. To dissect the impact of atmospheric pressure on stem cell state, we leveraged whole-transcriptome mRNA sequencing (mRNA-seq) on iPSCs cultured over time in variable atmospheric pressure and oxygen concentration. We cultured three unique human donor iPSC lines in 5% or 15% oxygen with either ambient (+ 0 PSI) or elevated (+ 2 PSI) atmospheric pressure and examined global gene expression changes by mRNA-seq and immunofluorescence staining for pluripotency and differentiation markers. Interestingly, and despite maintenance in medium supporting self-renewal, iPSCs cultured specifically in elevated atmospheric pressure and 15% oxygen for seven passages demonstrated morphologic and global gene expression changes consistent with stem cell differentiation and embryonic development, unlike their counterparts cultured in ambient atmospheric pressure or 5% oxygen. As early as passage three, iPSCs cultured in elevated atmospheric pressure and 15% oxygen exhibited a shift in metabolic gene expression and an up-regulation of genes involved in epigenetic regulation. To examine if atmospheric pressure promotes generation of lineage-specific cell types, we differentiated NSCs to both motor- and central nervous system (CNS)-type neurons using proteins / small molecules, and observed increased expression of neuronal maturation markers in response to pressure. We further show that atmospheric pressure can regulate hematopoietic differentiation of iPSCs through enrichment of a unique subset of CD34^+^CD43^low^CD45^high^ hematopoietic stem/progenitor cells (HSPCs) previously reported to exhibit greater potential for T-cell generation (Timmermans et al, 2009; Kennedy et al, 2012). In summary, our preliminary insights demonstrate that elevated atmospheric pressure during culture can enhance somatic cell reprogramming and stem cell differentiation workflows and offers a promising new strategy for generation of clinically important cell types.

## Results

### Elevated atmospheric pressure induces a 3-fold increase in somatic cell reprogramming to iPSCs across 10 unique human fibroblast donors

Previous findings highlight the regulatory role of hypoxia for maintenance of pluripotency and reprogramming of fibroblasts to iPSCs (Yoshida et al, 2009; Mathieu et al, 2013), therefore we investigated if elevated atmospheric pressure can further enhance reprogramming of human fibroblasts to iPSCs under hypoxic conditions. We transfected 10 unique human donor fibroblast lines with mRNA-based reprogramming factors (*OCT4, KLF4, SOX2, GLIS1, c-MYC* mRNAs) and found that transient exposure of fibroblasts to elevated atmospheric pressure (+ 5 PSI) and hypoxia (1% O_2_) for 24-hrs post-transfection followed by culture in 5% O_2_ and ambient atmospheric pressure resulted in ∼3-fold increase in reprogramming efficiency (Figures 1A-1C). Interestingly, static culture in 5% O_2_ with increasing atmospheric pressure had an inhibitory effect on reprogramming efficiency (Figure S1). Immunofluorescence staining for the pluripotency markers NANOG, POU5F1, SOX2, and SSEA-4 in iPSC lines established using this transient elevated pressure strategy did not indicate any loss of protein expression (Figure 1B). These results highlight a previously unknown role of atmospheric pressure during fibroblast re-programming that can provide for more robust iPSC generation without compromise to pluripotency.

**Figure 1.**
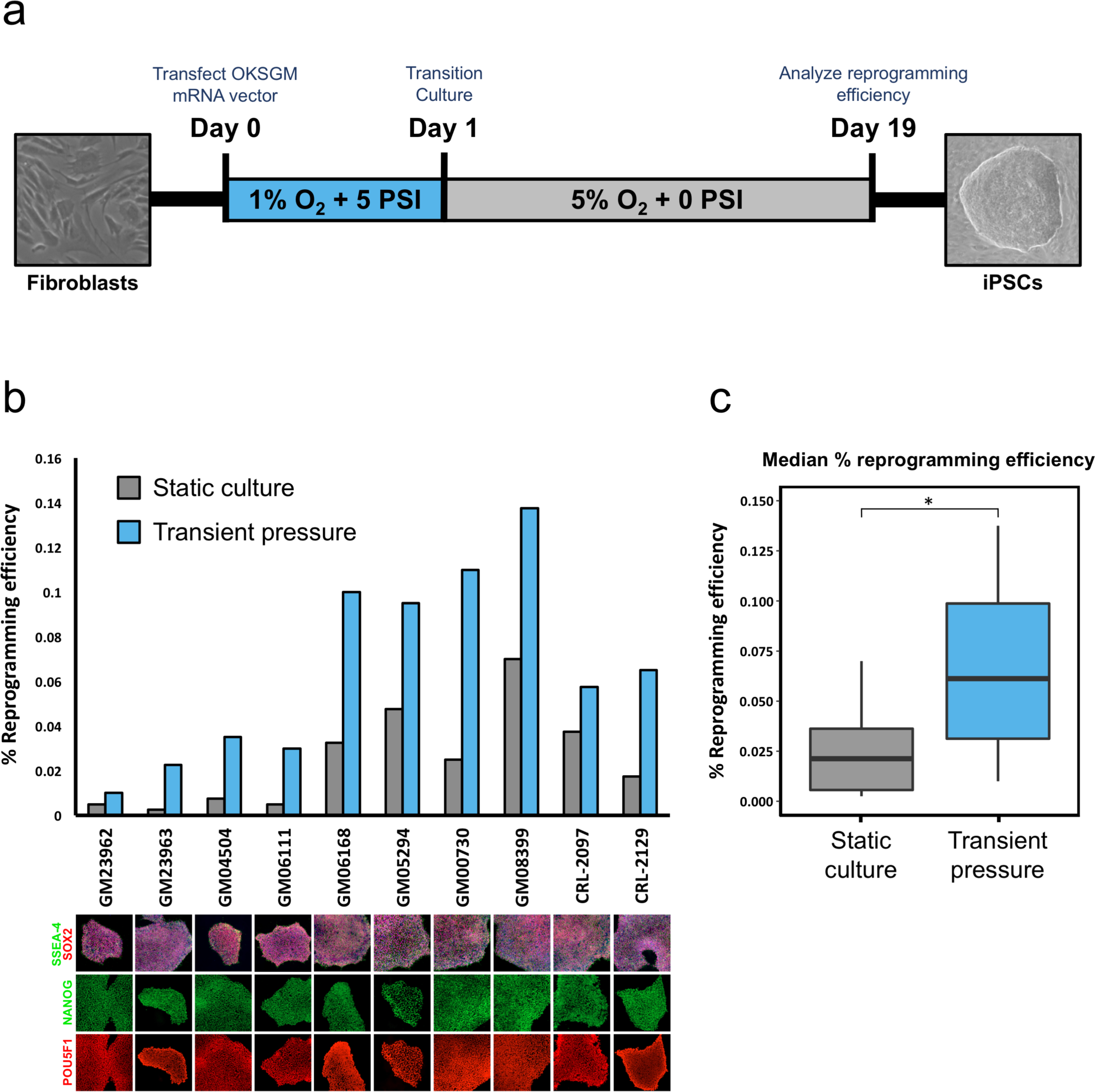
Elevated atmospheric pressure and hypoxia positively regulate re-programming of human fibroblasts to iPSCs. **(a)** Schematic of fibroblast re-programming strategy using transient elevated atmospheric pressure and hypoxia. **(b)** Re-programming of 10 unique human fibroblast lines using an mRNA reprogramming vector either in 5% O_2_ culture conditions for the entire experiment (static culture, gray) or transient exposure to 1% O_2_ + 5 PSI for 24 hr post-mRNA transfection followed by 5% O_2_ culture (transient pressure, blue) until colony counts were performed on day 19. Re-programming efficiency is calculated as the number of iPSC colonies / number of transfected fibroblasts. Representative immunofluorescence staining of each iPSC line at passage three for SSEA-4, SOX2, NANOG, and POU5F1 is shown below the column graph. **(c)** Boxplot showing the median % reprogramming efficiency for each culture condition across all 10 fibroblast lines. * = p-value < 0.05 student’s Ttest.

### Elevated atmospheric pressure induces altered expression of 56 genes associated with embryo development and cell differentiation

To investigate the role of atmospheric pressure and oxygen concentration during culture of iPSCs, we varied atmospheric pressure (+ 0 PSI and + 2 PSI, relative to ambient) and oxygen concentration (5% and 15% O_2_) and performed routine passaging under feeder-free conditions using three unique human donor iPSC lines from both fibroblast and cord blood sources; iPSC-1 and iPSC-2 from skin fibroblast, and iPSC-3 from cord blood CD34+ cells were used in this study. We cultured iPSCs feeder-free in mTesr1 medium and incubated in 15% O_2_ + 0 PSI or + 2 PSI, and 5% O_2_ + 0 PSI or + 2 PSI, and performed mRNA-seq and immunofluorescence profiling for pluripotency markers at the time-points illustrated in Figure 2A. By passage seven we observed 590 significant (false discovery rate (FDR) adjusted p-value < 0.05) differentially expressed genes unique in 15% O_2_ + 2 PSI relative to 15% O_2_ + 0 PSI, and 348 similarly differentially expressed genes unique in 5% O_2_ + 2 PSI relative to 5% O_2_ + 0 PSI, with only 56 genes overlapping between these two groups (Figures 2B & 2C). Of these 56 “pressure-specific” overlapping genes significantly differentially expressed in elevated pressure in both 15% and 5% O_2,_ many play a role in embryo development, cell differentiation, ECM composition, and metabolism (Figure 2C). These observations suggest that elevated atmospheric pressure influences distinct sets of genes synergistically with differential oxygen concentration as evidenced by the unique expression profile of iPSCs cultured in 15% O_2_ + 2 PSI relative to 5% O_2_ + 2 PSI (Figures 2B, 2C and S3A).

**Figure 2.**
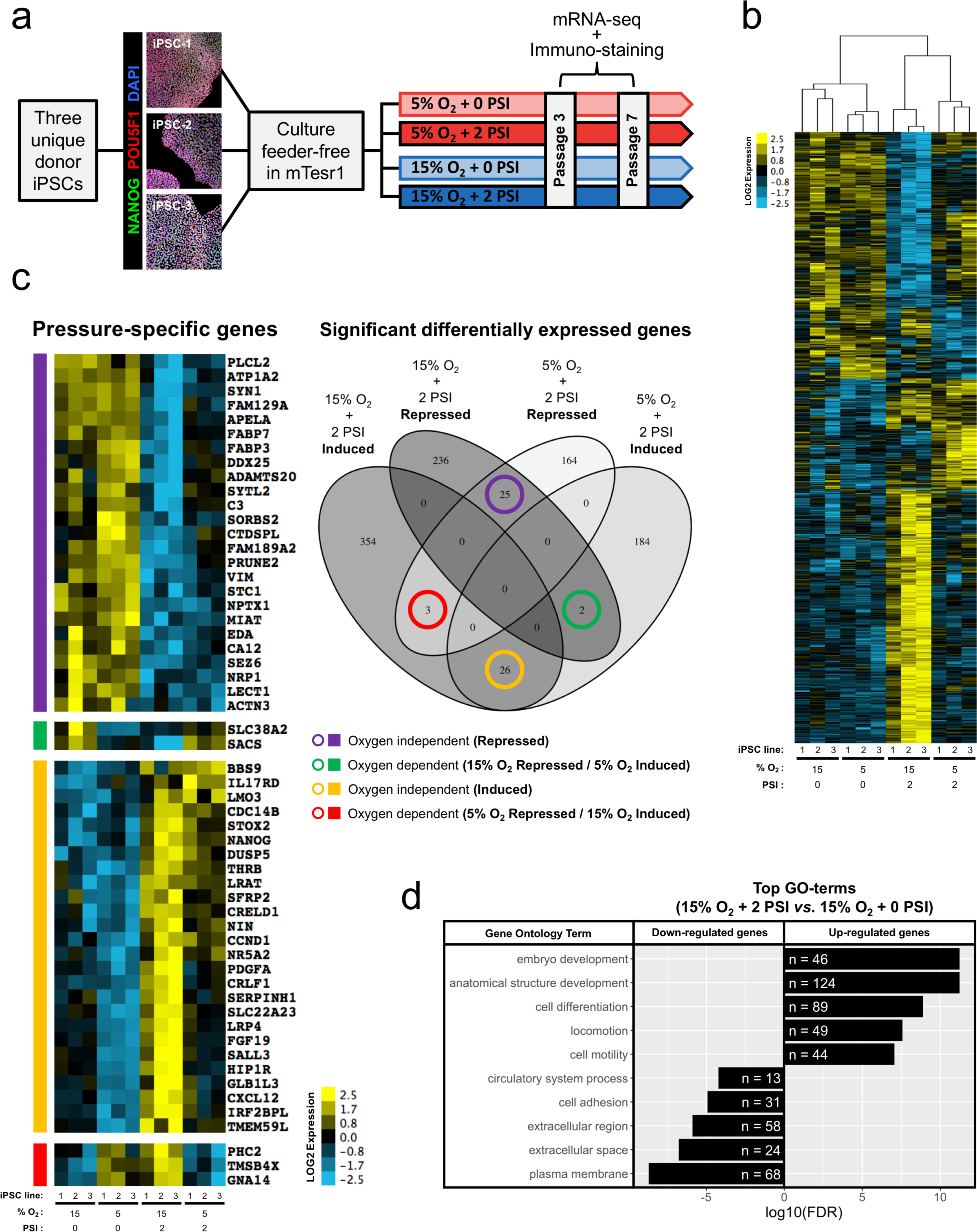
Culture of iPSCs in elevated atmospheric pressure induces transcriptional changes associated with epigenetic regulation and differentiation. **(a)** Schematic of experimental design. **(b)** Unsupervised hierarchical clustering of cumulative differentially expressed genes (763 total) by whole-transcriptome RNA-seq at passage seven in the indicated O_2_ and PSI with adjusted (FDR) p-value < 0.05 between 15% O_2_ + 2 PSI *vs.* 15% O_2_ + 0 PSI, and 5% O_2_ + 2 PSI *vs.* 5% O_2_ + 0 PSI using DEseq2. n = 3 independent donor iPSC lines, 2 from human fibroblast (iPSC lines 1 and 2) and 1 from human CD34+ cord blood (iPSC line 3) cells. Data in heatmap is displayed as log2 transformed gene expression. **(c)** Venn diagram of significant differentially expressed genes from **(b)** showing overlap between analysis groups and heatmap showing overlapping genes that are significantly differentially expressed in response to elevated pressure in both 5% and 15% O_2_. **(d)** The top 5 GO terms for both up-regulated and down-regulated differentially expressed genes between 15% O_2_ + 2 PSI *vs.* 15% O_2_ + 0 PSI at passage seven. Gene ontology enrichment is ranked according to adjusted p-value (FDR) and displayed on log10 scale. The number of significant differentially expressed genes for each GO term is displayed inside the bar graph.

### Elevated atmospheric pressure increases pluripotency gene transcription and retains protein expression of POU5F1, NANOG, and SOX2 largely independent of hypoxia

To further define the differentiation state of iPSCs cultured to passage seven in 15% O_2_ + 2 PSI we examined the expression levels of germ layer commitment and pluripotency-regulating genes. Interestingly, the protein expression of pluripotency-regulating genes POU5F1, NANOG, and SOX2 are largely retained in iPSCs in 15% O_2_ + 2 PSI, except in areas of the cultures that are distinct from the tight colony formations typical of un-differentiated iPSCs (Figure 3C). Phase contrast images of iPSCs cultured in 15% O_2_ + 2 PSI indicate a larger cytoplasm-to-nucleus ratio relative to ambient pressure controls or iPSCs cultured in 5% O_2_, a feature common in differentiated pluripotent stem cells (Figure 3A). Surprisingly, expression levels of several pluripotency-regulating genes are significantly increased in iPSCs in 15% O_2_ + 2 PSI relative to 15% O_2_ + 0 PSI, including *GDF3*, *MMP2*, *NANOG*, *NR5A2*, and *POU5F1*, whereas in 5% O_2_ only *NANOG* and *NR5A2* reached statistical significance (Figure 3B). Despite the increased expression of pluripotency-regulating genes, canonical germ layer commitment markers are significantly up-regulated in iPSCs in 15% O_2_ + 2 PSI, with a predominance of endodermal lineage genes (Figure 3B).

**Figure 3.**
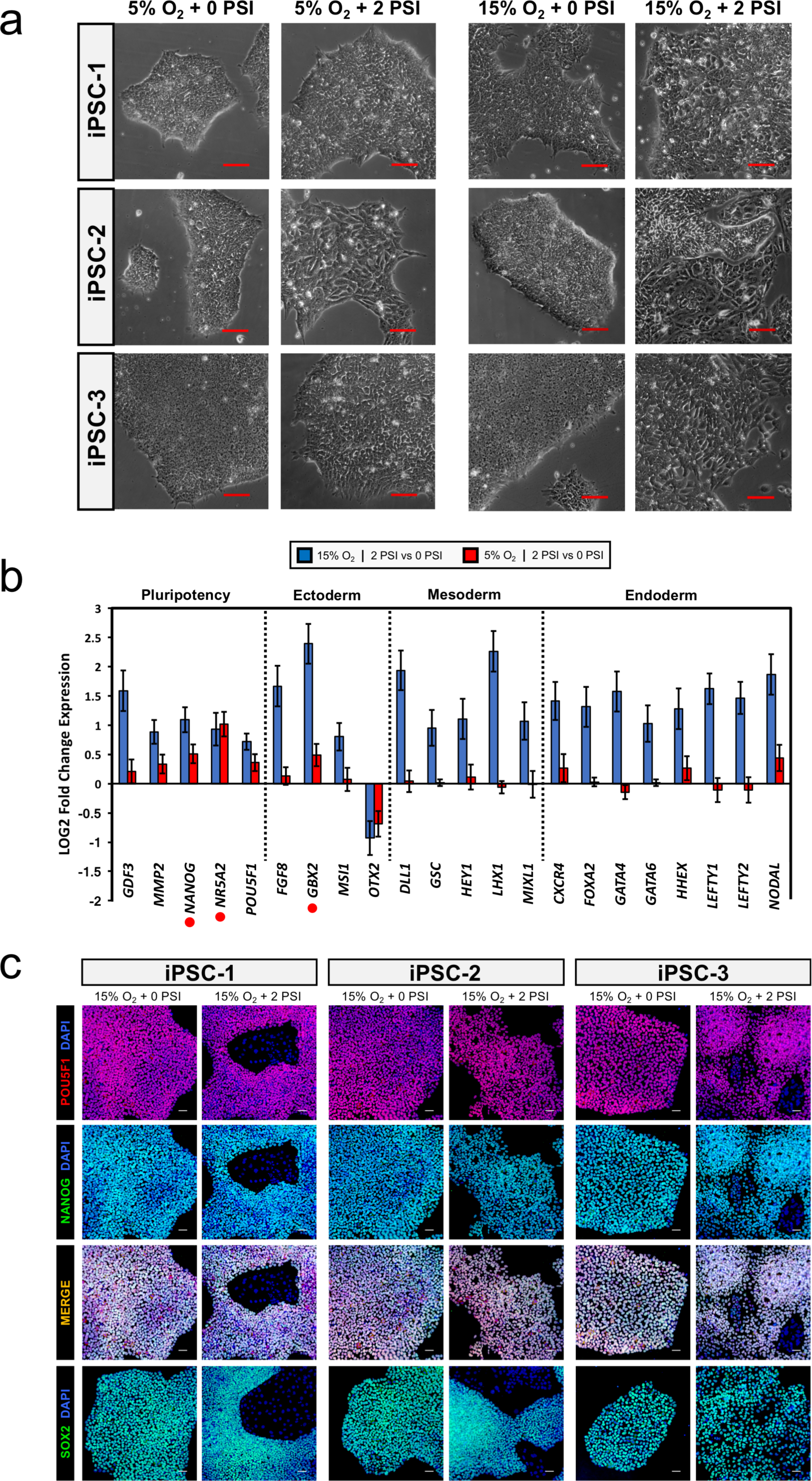
iPSCs cultured long-term in elevated atmospheric retain pluripotency with concomitant increases in differentiation markers and loss of morphology. **(a)** Representative phase contrast images of passage seven iPSCs in the indicated culture conditions. Scale bars are 200 um.

Moreover, gene ontology enrichment analysis of iPSCs in 15% O_2_ + 2 PSI identified pathways related to embryo development and cell differentiation with up-regulated genes, and pathways related to cell adhesion, extracellular space/region, and plasma membrane with down-regulated genes (Figure 2D & Supplementary data). Conversely, gene ontology enrichment analysis of iPSCs in 5% O_2_ + 2 PSI identified cell differentiation pathway genes as down-regulated at both passage three and seven, indicating a strong inhibitory effect of lower oxygen in conjunction with elevated pressure on these pathway gene members (Figure S2A & Supplementary data). In summary, iPSCs cultured in 15% O_2_ + 2 PSI largely retain expression of POU5F1, NANOG, and SOX2 with concomitant increases in subsets of pluripotency and germ layer commitment markers.

### Up-regulation of genes associated with primed- and naïve-state pluripotency and NODAL/LEFTY signaling pathway in iPSCs cultured short-term in elevated atmospheric pressure

To elucidate the genetic perturbations that precede the differentiation phenotype observed at passage seven in iPSCs cultured in 15% O_2_ + 2 PSI, we analyzed mRNA-seq data of iPSCs at passage three for differentially expressed genes in 15% O_2_ + 2 PSI *vs.* 15% O_2_ + 0 PSI. We observed an approximately 4-fold increase in the number of significant (FDR adjusted p-value < 0.05) differentially expressed genes in 15% O_2_ + 2 PSI *vs.* 15% O_2_ + 0 PSI (1473 total) compared to 5% O_2_ + 2 PSI *vs.* 5% O_2_ + 0 PSI (358 total), with 117 overlapping genes between the two groups (Figures 4A & 4F, Figure S4). To better characterize the pluripotent state of iPSCs cultured short-term in elevated atmospheric pressure, we examined differential expression of genes involved in naïve- and primed-state pluripotency. Surprisingly, we observed significant increases in expression levels of both naïve- and primed-state pluripotency markers in iPSCs in 15% O_2_ + 2 PSI (Figure 4B), suggesting an intermediary pluripotent state. The mRNA expression level of *NANOG* is significantly higher in both 15% O_2_ + 2 PSI and 5% O_2_ + 2 PSI relative to ambient atmospheric pressure controls (Figure 4B). NANOG protein expression is progressively up-regulated during murine embryonic development and remains high in the inner cell mass population that specifies formation of the epiblast (Komatsu et al, 2015). Similarly, *DPPA3*, a gene enriched in human pre-implantation epiblast cells (Chan et al, 2013) is also up-regulated in elevated pressure, but specifically in 15% O_2_ (Figure 4B).

**Figure 4.**
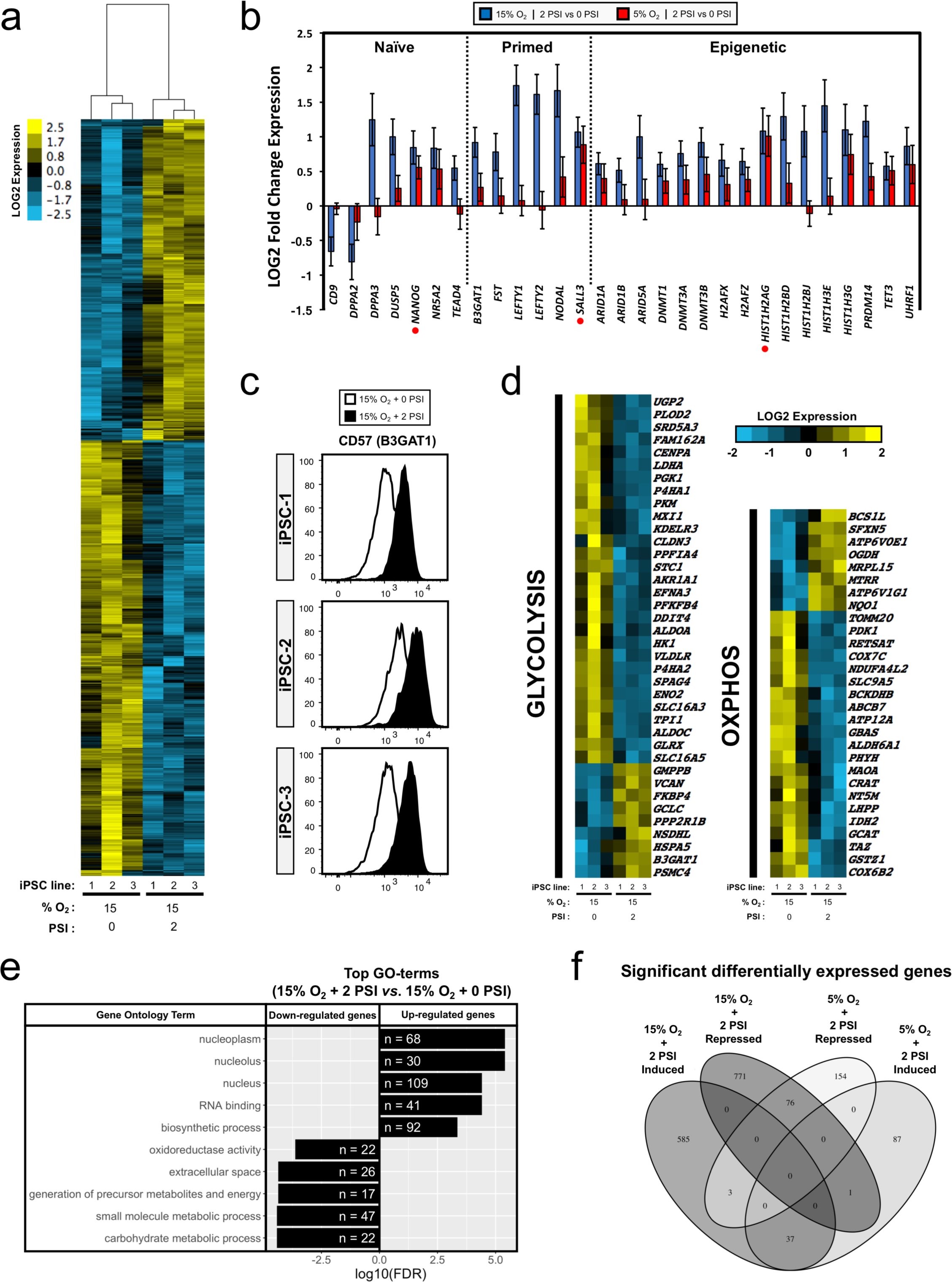
A shift in metabolic and epigenetic gene expression signature is observed in iPSCs cultured short-term in elevated atmospheric pressure. **(a)** Unsupervised hierarchical clustering of differentially expressed genes (1473 total) by whole-transcriptome mRNA-seq at passage three with FDR adjusted p-value < 0.05 between 15% O_2_ + 2 PSI *vs*. 15% O_2_ + 0 PSI. n = 3 independent donor iPSC lines as described in (**2a**). **(b)** Naïve/Primed pluripotency-associated and epigenetic-regulator gene expression from DEseq2 represented as log2 fold changes between both 15% O_2_ + 2 PSI *vs.* 15% O_2_ + 0 PSI (blue bars) and 5% O_2_ + 2 PSI *vs.* 5% O_2_ + 0 PSI (red bars) iPSC lines. Error bars represent standard error of log2 fold change. All 15% O_2_ + 2 PSI *vs.* 15% O_2_ + 0 PSI (blue bars) have FDR adjusted p-value < 0.05, and those gene names with an orange dot are also significant for 5% O_2_ + 2 PSI *vs.* 5% O_2_ + 0 PSI (red bars). n = 3 as described in **(2a)**. **(c)** Flow cytometry analysis of passage three iPSCs stained for the primed-state pluripotency associated surface marker CD57 (B3GAT1). **(d)** Glycolysis and oxidative phosphorylation (OXPHOS) pathway-related genes from Hallmark, GO, and Kegg gene ontology databases that are differentially expressed between 15% O_2_ + 2 PSI *vs*. 15% O_2_ + 0 PSI at passage three with FDR adjusted p-value < 0.05. Un-supervised hierarchical clustering of genes / samples shown as scaled log2 fold change across rows (genes). **(e)** The top 5 GO terms for both up-regulated and down-regulated differentially expressed genes between 15% O_2_ + 2 PSI *vs*. 15% O_2_ + 0 PSI at passage three. Gene ontology enrichment ranked according to FDR adjusted p-value and displayed on log10 scale. The number of significant differentially expressed genes for each GO term is displayed in the bar graph. **(f)** Venn diagram of significant differentially expressed genes from (**a**) and additionally, differentially expressed genes between 5% O_2_ + 0 PSI *vs*. 5% O_2_ + 2 PSI showing overlap between groups.

The primed-state pluripotency-associated DNA methyltransferase family genes *DNMT3A* and *DNMT3B* are significantly up-regulated in 15% O_2_ + 2 PSI relative to 15% O_2_ + 0 PSI (Figure 4B), which are reported to increase in expression during transition from pre- to post-implantation epiblast stage of mammalian embryogenesis (Boroviak et al, 2015). We next evaluated the protein expression level of the primed-state pluripotency-associated surface marker B3GAT1 (Collier et al, 2017) by flow cytometry in iPSCs in 15% O_2_ + 2 PSI and observed an approximately 0.5 log increase in fluorescence intensity relative to 15% O_2_ + 0 PSI across all three iPSC lines (Figure 4C), with only a modest increase in 5% O_2_ + 2 PSI (Figure S2A). Furthermore, the transforming growth factor beta family member *NODAL*, as well as its targets *LEFTY1* and *LEFTY2*, are highly up-regulated in iPSCs cultured in 15% O_2_ + 2 PSI. These NODAL-signaling pathway genes are known to be up-regulated during epiblast priming in mammalian embryogenesis or during transition from naïve state pluripotency in murine embryonic stem cells (ESCs) cultured in-vitro (Figure 4B) (Boroviak et al, 2015, Mulas et al, 2016). In summary, these observations suggest that iPSCs cultured to passage three in 15% O_2_ + 2 PSI are in a pluripotent state with transcriptional features in common with the epiblast-stage of the developing mammalian embryo.

### Elevated atmospheric pressure promotes iPSC differentiation through epigenetic-regulatory gene expression changes independent of hypoxia

The epigenetic modifications that occur during both in vitro pluripotent stem cell differentiation and in developing pre-implantation embryos consist of a progressive genome-wide establishment of methylation patterning and histone modifications (Wu et al, 2006; Hawkins et al, 2010; Gifford et al, 2013). DNA methylation during differentiation of pluripotent stem cells and in early embryogenesis is facilitated by DNA methyltransferases DNMT1, DNMT3A, and DNMT3B, along with co-factors such as UHRF1 (Boland et al, 2014) and the TET-family dioxygenase TET3 (Kang et al, 2015). The differential gene expression levels for these DNA methylation-regulating enzymes are significantly up-regulated in iPSCs cultured to passage three in 15% O_2_ + 2 PSI *vs.* 15% O_2_ + 0 PSI, as are the chromatin remodeling genes *ARID1A*, *ARID1B*, and *ARID5A* (Figure 4B), of which ARID1A is shown to be essential for murine ESC differentiation (Baba et al, 2011; Gao et al, 2008). Interestingly, increased expression of genes encoding histone subunits are observed in 15% O_2_ + 2 PSI (Figure 4B), possibly to meet the demand of progressive methylation in the DNA landscape. Moreover, gene ontology enrichment analysis revealed that pathways involved in nuclear processes and RNA-binding contain genes significantly up-regulated in 15% O_2_ + 2 PSI (Figure 4E, supplementary data), suggesting increased transcriptional and/or chromosomal activity in these iPSCs. These findings provide further evidence that iPSCs in 15% O_2_ + 2 PSI are transitioning to a more differentiated state.

### Elevated atmospheric pressure down-regulates glycolysis and OXPHOS metabolic pathways independent of hypoxia

Gene ontology enrichment analysis of iPSCs cultured to passage three in 15% O_2_ + 2 PSI indicates pathways governing carbohydrate and small molecule metabolic processes, as well as oxidoreductase activity, contain genes that are significantly down-regulated (Figure 4E). Additionally, differential gene expression analysis of 15% O_2_ + 2 PSI *vs.* 15% O_2_ + 0 PSI revealed that 38 genes involved in glycolysis and 29 involved in oxidative phosphorylation (OXPHOS) are significantly differentially expressed (FDR adjusted p-value < 0.05), with the majority down-regulated (62% and 69%, respectively) in response to elevated pressure (Figure 4D). The prevalent down-regulation of glycolysis- and OXPHOS-related genes in 15% O_2_ + 2 PSI may be a result of differentiation, as a previous study demonstrated decreased expression of TCA-cycle and glycolysis-related genes in fibroblasts cells compared to counterpart iPSC lines (Varum et al, 2011). Only 7 genes involved in glycolysis and 6 involved in OXPHOS are significantly differentially expressed in 5% O_2_ + 2 PSI *vs.* 5% O_2_ + 0 PSI (Figure S2B), indicating a contribution of higher oxygen to the shift in metabolic gene expression (Figure S2C). In summary, iPSCs cultured in 15% O_2_ + 2 PSI exhibit a general down-regulation of glycolysis- and OXPHOS-related gene expression.

### Neural-ectoderm markers PAX6 and NES are increased during directed-differentiation of iPSCs in response to elevated atmospheric pressure

Physical force-mediated regulation of neuronal differentiation in adult stem cells has been achieved through modulation of substrate rigidity (Engler et al, 2006; Saha et al, 2008; Leipzig et al, 2009). To investigate if elevated atmospheric pressure exerts a similar regulatory influence we cultured iPSCs in neural induction medium in varying oxygen (5% and 15% O_2_) and atmospheric pressure (+ 0 PSI and + 2 PSI) and evaluated the expression of the neural-ectoderm markers PAX6 and NES by immuno-fluorescence imaging and flow cytometry. After 6 days of differentiation, iPSCs cultured in 5% O_2_ + 2 PSI exhibited 93% of cells co-expressing PAX6 and NES and a ∼2-fold increase in mean fluorescence intensity of protein staining by flow cytometry relative to 5% O_2_ + 0 PSI (Figure 5A). Surprisingly, in the absence of the neutralizing factor NOGGIN in induction medium, iPSCs cultured in 5% O_2_ + 2 PSI exhibited 90% of cells co-expressing PAX6 and NES, approximately 5-fold greater than ambient pressure controls or iPSCs cultured in 15% O_2_ + 2 PSI (Figure 5A), demonstrating that elevated atmospheric pressure and hypoxia are significant drivers of neuronal differentiation.

**Figure 5.**
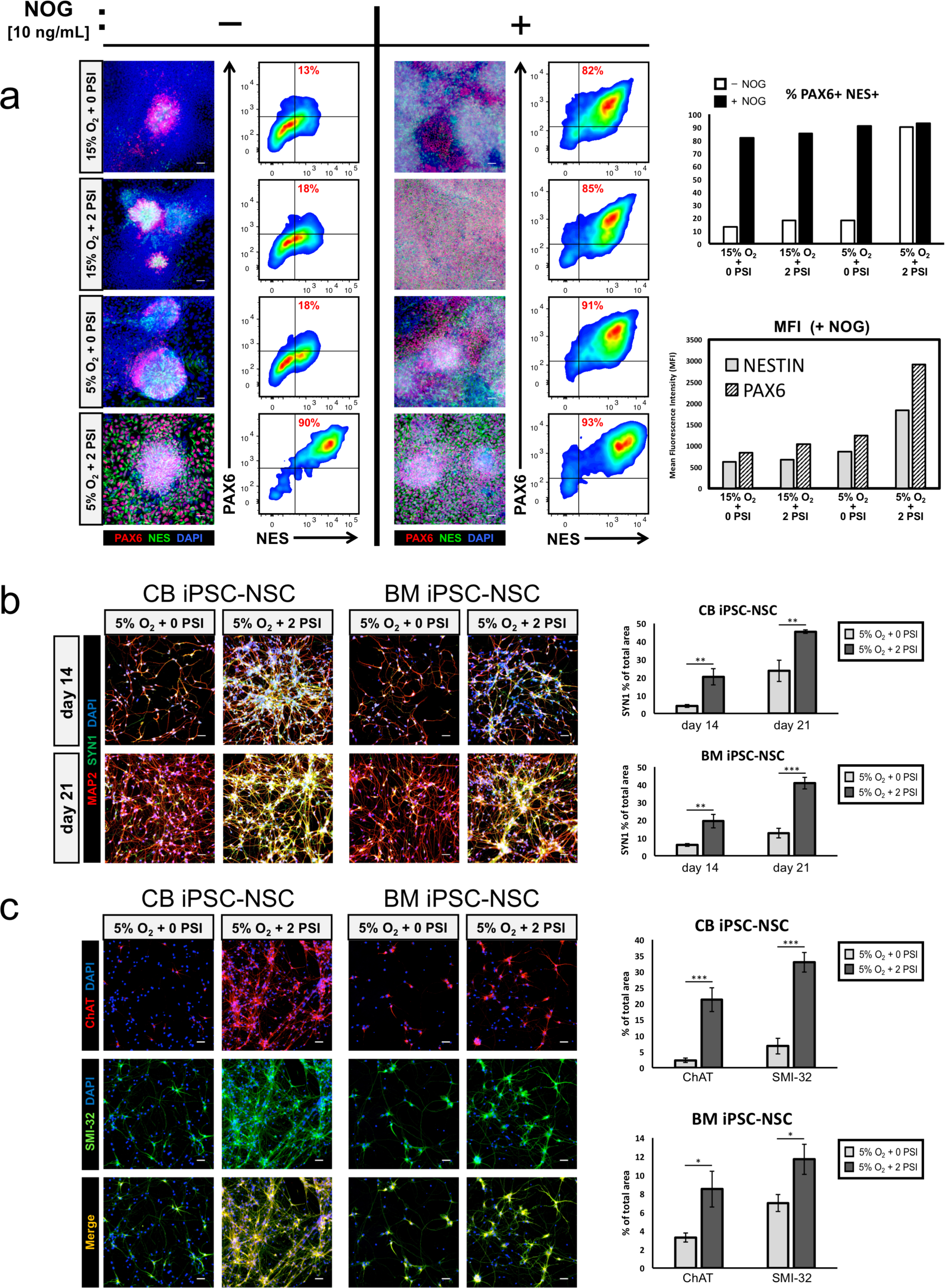
Elevated atmospheric pressure can regulate iPSC differentiation to ectodermal and mesodermal lineages and enhance re-programming of somatic cells. **(a)** Representative immunofluorescence images and parallel flow cytometry analysis of iPSC-2 line differentiated for 6 days in medium containing 10 ng/mL recombinant LIF with and without 10 ng/mL NOG and stained for PAX6 and NES. Top column graph shows % PAX6 and NES double positive cells from flow cytometry data. Bottom column graph shows geometric mean fluorescence of PAX6 and NES staining from flow cytometry data. Scale bar = 50 um. **(b)** CNS-type neuronal differentiation from cord blood derived iPSC neural stem cells (CB iPSC-NSC) and bone marrow derived iPSC neural stem cells (BM iPSC-NSC) stained for both MAP2 (red) and SYN1 (green) day 14 and 21 of differentiation. Scale bars = 50 uM. Quantification of the mature neuronal marker SYN1 is displayed in the column graphs to the right. **(c)** Representative immunofluorescence images of day 31 motor neurons differentiated using the same source NSC lines as in **(b)** stained for Choline Acetyltransferase (ChAT, in red) and Neurofilament-H (SMI-32, in green). Scale bars = 50 um. Quantification for ChAT and SMI-32 is displayed in the column graphs to the right as % of total area of staining. For both **(b)** and **(c)**, n = 3 images quantified per condition. Error bars are standard error of the mean. * = p-value < 0.05, ** = < 0.01 student’s Ttest.

### Elevated atmospheric pressure and hypoxia induces ChAT and NF-H in motor-neurons and SYN1 in CNS-type neurons

Based on our neural induction results and previous reports showing that low oxygen promotes differentiation and maturation of neuronal cell types (Xie et al, 2014; Yasui et al, 2017), we next asked if elevated pressure acts synergistically with 5% O_2_ to generate terminally differentiated neurons. We differentiated 2 unique NSC lines, one derived from a cord-blood sourced iPSC line (CB iPSC-NSC) and one from a bone marrow sourced iPSC line (BM iPSC-NSC), to both motor and CNS-type neurons in the presence of 5% O_2_ + 2 PSI. Following 31 days of NSC differentiation to motor neurons, we stained the cells for the mature motor neuron markers choline acetyltransferase (ChAT) and neurofilament-H (SMI-32) and observed a significant increase in expression of both markers in 5% O_2_ + 2 PSI (Figure 5C). We observed a similar trend upon differentiation of NSCs to CNS-type neurons; immunofluorescence staining for MAP2 and the mature neuronal marker SYN1 at days 14 and 21 of differentiation showed a significant increase in SYN1 expression in 5% O_2_ + 2 PSI (Figure 5B). Taken together, these results show that elevated atmospheric pressure when combined with hypoxia, positively regulates differentiation and maturation of neuronal lineages.

### Elevated atmospheric pressure enriches for a CD43^low^CD45^high^ hematopoietic lineage subset associated with T-lymphocyte commitment

We next evaluate the influence of elevated atmospheric pressure on mesoderm lineage commitment. Physical force in the form of shear stress has been leveraged to enhance ESC differentiation to endothelial progenitor and hematopoietic lineages (Wolfe and Ahsan, 2013), and a combination of osteoblasts and intermittent hydrostatic pressure has been used to show an influence on HSPC migration (Kim et al, 2019; Lee et al, 2020), however no studies have implemented hydrostatic pressure to generate HSPCs from pluripotent stem cells on their own. We differentiated iPSCs to HSPCs expressing CD34, CD43, and CD45 using a bi-phasic oxygen concentration protocol to simulate in-vivo HSPC development. The hematopoietic stem cell niche is progressively lower in oxygen at greater distances from the bone marrow sinuses, where the most primitive stem cells reside (Chow et al, 2001). We found that transitioning from 3% O_2_ + 2 PSI to 15% O_2_ + 2 PSI enriched a sub-population of cells expressing CD34^+^CD43^low^CD45^high^ compared to ambient atmospheric pressure controls (Figures 6A & 6B). Sorting and further differentiation of this subset of CD34^+^CD43^low^CD45^high^-expressing HSPCs has been reported to possess lineage bias for T-lymphocyte commitment (Timmermans et al, 2009, Kennedy et al, 2012), suggesting that T-lymphocyte generation from iPSCs can be enhanced by implementation of elevated atmospheric pressure to serve as a potent differentiation cue.

**Figure 6.**
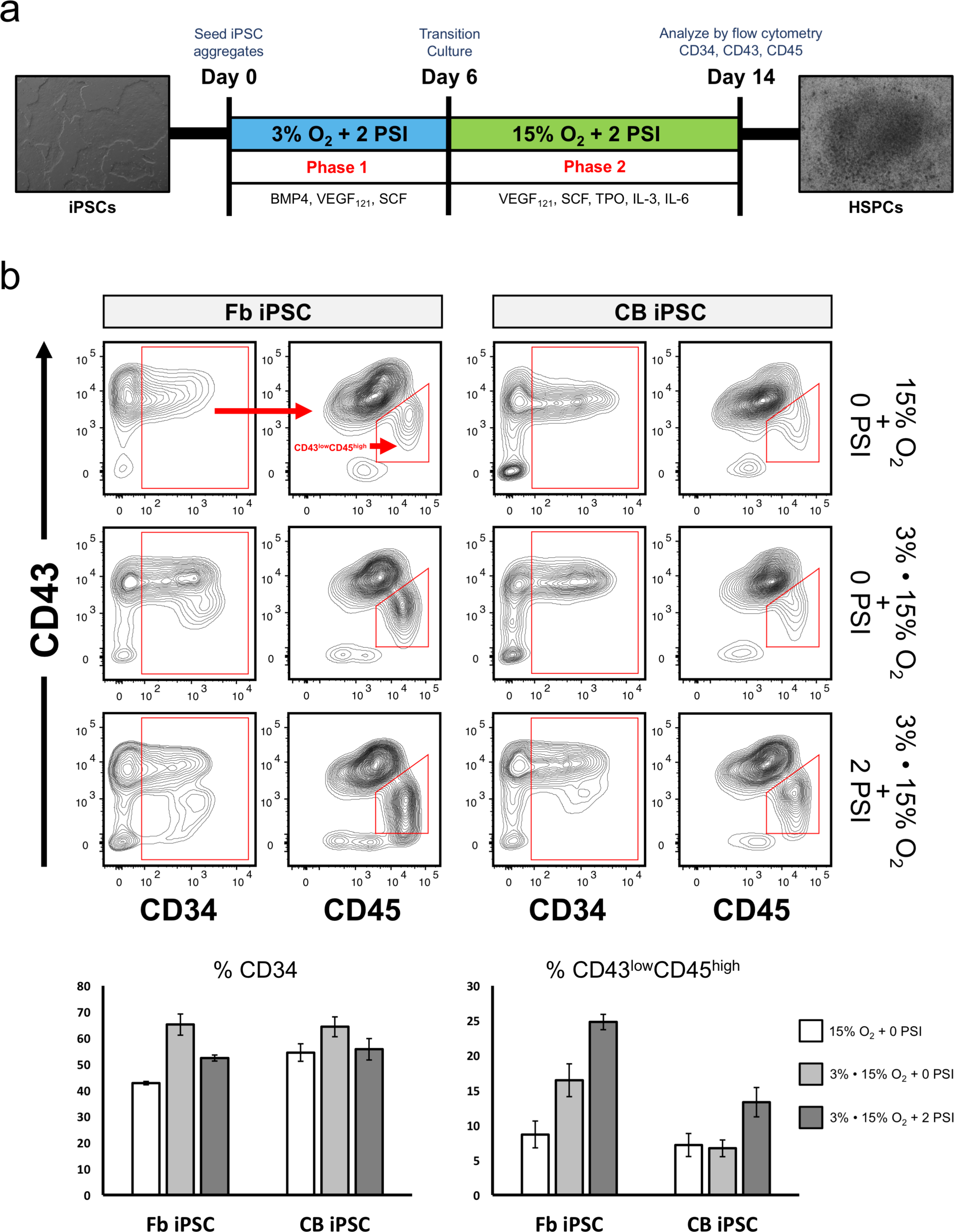
Elevated atmospheric pressure regulates iPSC differentiation to a specific subset of CD34^+^CD43^low^CD45^high^ HSPCs. **(a)** Schematic of hematopoietic stem cell differentiation protocol showing incubation strategy to enrich for CD34^+^CD43^low^CD45^high^ subset. **(b)** Hematopoietic stem cell differentiation from a fibroblast-derived iPSC line (Fb iPSC) and cord blood iPSC line (CB iPSC) stained for CD34, CD43, and CD45 and analyzed by flow cytometry. Conditions denoted with 3% • 15% O_2_ were transitioned from 3% to 15% O_2_ at day 6 of differentiation and allowed to culture for a further 8 days before analysis. The column graph below shows the quantification of the percent CD34+ (left) and CD34^+^CD43^low^CD45^high^ (right) population indicated by red gates in the plots. Error bars are standard error of the mean. **(b)** Pluripotency and germ layer differentiation marker gene expression from DEseq2 mRNA-seq analysis between both 15% O_2_ + 2 PSI *vs.* 15% O_2_ + 0 PSI (blue bars) and 5% O_2_ + 2 PSI *vs*. 5% O_2_ + 0 PSI (red bars) iPSC lines. Error bars represent standard error of log2 fold change. Adjusted p-value (FDR) < 0.05 for all 15% O_2_ + 2 PSI *vs.* 15% O_2_ + 0 PSI differentially expressed genes and those gene names with an orange dot are also significant for 5% O_2_ + 2 PSI *vs.* 5% O_2_ + 0 PSI (red bars). n = 3 independent iPSC lines as described in (**2a**). **(c)** Representative immunofluorescence images of passage seven iPSCs showing expression of POU5F1, NANOG, and SOX2 in the indicated culture conditions. Scale bars are 50 um.

## Discussion

Physical forces can regulate differentiation of both adult and pluripotent stem cells in-vivo and in-vitro. Here we demonstrate a novel bioreactor that can apply physical force using atmospheric pressure to enhance somatic cell reprogramming and induce differentiation of both iPSCs and NSCs. Elevated atmospheric pressure (+ 2 PSI) in conjunction with 15% oxygen during extended culture of iPSCs resulted in a differentiation phenotype characterized by increased germ-layer commitment and pluripotency marker expression and enrichment of genes involved in embryo development and cell differentiation gene ontology pathways. iPSCs cultured short-term in 15% O_2_ + 2 PSI exhibit up-regulation of epigenetic-regulatory gene expression and a general down-regulation of metabolic gene expression. Furthermore, we demonstrated an enhancement in differentiation of iPSCs to neural-ectoderm and NSCs to motor and CNS-type neurons by culture in elevated atmospheric pressure and 5% oxygen. Lastly, enrichment of a hematopoietic progenitor CD34^+^CD43^low^CD45^high^ population with T-lymphocyte lineage bias was achieved by simultaneous culture in elevated atmospheric pressure and transition from low to high oxygen. These findings shed light on a previously unknown role for atmospheric pressure as a physical force during iPSC culture and stem cell differentiation.

How does elevated atmospheric pressure act to induce stem cell differentiation? Fluid-pressure measurements within the mouse blastocyst reveals increased pressure over the course of development, thereby generating a physical force input and a potential cue for embryo morphogenesis (Wang et al, 2018). In vitro application of physical force can influence stem cell differentiation through activation of mechano-transduction pathways (Northey et al, 2017). For example, transmembrane integrins, stretch activated ion channels, and focal adhesion proteins are all implicated in physical force-directed cell signaling to trigger both phenotypic and functional responses, including cell differentiation (Formigli et al, 2007; Chistiakov et al, 2017).

Additionally, cross-talk between mechano-sensing pathways and cellular metabolism are known to exist in cancer cells (Boulter et al, 2018), thus it is possible atmospheric pressure is triggering a similar response in iPSCs and NSCs given our observation that metabolic gene expression is significantly altered in elevated atmospheric pressure.

Metabolism is known to regulate differentiation states of pluripotent stem cells (Prigione and Adjaye, 2010), however it is unclear if our observations of metabolic gene expression changes are a direct effect of elevated atmospheric pressure or rather, a consequence of the differentiation phenotype. Future experiments employing a transient exposure or dose-response of elevated atmospheric pressure would enable interrogation of mechano-sensing pathways.

Elevated atmospheric pressure-mediated physical force could be exerting its regulatory influence on stem cell state by a similar mechanism to other well-characterized forms of force that are currently utilized in cell culture. For example, physical force in the form of substrate rigidity has been leveraged towards in-vitro stem cell differentiation by modulating ECM composition and stiffness of the culture surface (Engler et al, 2006; Saha et al, 2008; Leipzig et al, 2009). Specifically, Engler et al showed using Collagen-I coated gel substrates of tunable rigidity that lineage specification during mesenchymal stem cell differentiation is mediated through mechano-sensing of physical force via ECM focal-adhesion complexes. Substrate viscoelasticity has also been shown to influence lineage commitment through modulation of ECM protein composition, with measurable changes in viscoelasticity depending on the concentration and combination of Collagen-1/4, Laminin, and Heparan Sulfate (Raghavan and Bitar, 2014). These findings demonstrate that external physical cues generated by substrate rigidity and/or manipulation of ECM composition in the microenvironment can direct stem cell differentiation.

Another attractive method to generate physical force in stem cell culture is via three-dimensional cellular models to mimic cell-to-cell interactions (Eiraku and Sasai, 2012; Zhong et al, 2014; Yin et al, 2016). These three-dimensional cellular models exploit the stem cell’s innate ability to differentiate into adult cell types and self-organize into the appropriate tissue architecture corresponding to the lineage identity of the stem cell (Murrow et al, 2017). The orchestration of this self-assembly in-vivo not only requires cell-to-cell interactions via adhesion molecules (Runswick et al, 2001; Chanson et al, 2011), but likely some contribution from physical forces to act as differentiation cues, as in-vitro differentiation is heavily influenced by ECM spatial arrangement and composition (Yin et al, 2016). The recapitulation of three-dimensional cell-to-cell interactions is a promising strategy for stem cell differentiation, however these models still generally lack the contribution of physical force by neighboring tissues that exists in-vivo. Elevated atmospheric pressure could simulate physical force exerted by the surrounding tissue microenvironment and combination with these three-dimensional models warrants future investigation.

Lastly, fluid mechanical force or “shear stress,” is common to cells residing in or surrounding the vascular system in-vivo and plays a significant role in cell differentiation. For example, vascular smooth muscle cells and endothelial cells that make up the lining of blood vessels are subject to variable degrees of shear stress due to haemodynamic forces, which in turn elicit phenotypic and functional responses to maintain homeostasis of the circulatory system (Chistiakov et al, 2017). During in vitro embryonic stem cell differentiation, fluid-mechanical force in the form of cyclic strain or shear stress can induce vascular smooth muscle cell and cardiomyocyte differentiation, respectively (Shimizu et al, 2008; Correia et al, 2014). Although haemodynamic forces are inherently variable relative to those found in most adult tissues in the human body, which are by comparison mechanically static, tissue homeostasis is likely maintained by tissue-specific tensional homeostasis (Northey et al, 2017). By this reasoning, physical force should be applied in a manner that recapitulates development of the desired tissue type for in vitro stem cell differentiation protocols.

The observation of a synergistic effect of elevated atmospheric pressure and 15% O_2_, but not 5% O_2_, on iPSC differentiation under standard stem cell culture conditions is unclear, but could be explained in part by the known positive regulation of pluripotency by hypoxia (Yoshida et al, 2009; Mathieu et al, 2013). For example, hypoxia inducible factors HIF1A and HIF1B are necessary for metabolic shift from oxidative phosphorylation to glycolysis during reprogramming of somatic cells to iPSCs (Mathieu et al, 2014) and HIF2A is shown to activate Oct4 (mouse homolog of POU5F1) in mouse embryonic stem cells (Covello et al, 2006). Therefore, the influence of elevated atmospheric pressure on iPSC differentiation may be negated or diminished by culture in 5% O_2_, which is sufficiently hypoxic for HIF protein stabilization.

Conversely, 5% O_2_ and elevated atmospheric pressure was demonstrated to be optimal for directed-differentiation of iPSCs to neural-ectoderm and NSCs to motor and CNS-type neurons. The enhancement of neural induction observed in 5% O_2_ is not surprising given that physiologic oxygen concentration in the developing mammalian brain is between 1 to 5% (Studer et al, 2000). Moreover, the positive influence of physical force on neuronal differentiation is well-established (Engler et al, 2006; Saha et al, 2008; Leipzig et al, 2009), therefore the combined contribution of 5% oxygen and elevated atmospheric pressure in our studies highlights the importance of mimicking the relevant tissue micro-environment for in vitro stem cell differentiation. This concept was further demonstrated for HSPC differentiation using a culturing protocol consisting of a combination of bi-phasic oxygen concentration and elevated atmospheric pressure to mimic hematopoietic development in the bone marrow niche. In summary, we have demonstrated a novel technology to improve stem cell generation and differentiation by modulating oxygen and atmospheric pressure simultaneously in the culture microenvironment. Importantly, this technology can be leveraged with existing stem cell differentiation protocols to improve efficiency of generating mature cell types for translational studies and clinical applications.

## Experimental Procedures

### Fibroblast re-programming

Human fibroblasts lines were obtained from the Coriell Institute or ATCC (see below table containing line information) and cultured in Eagle’s Minimum Essential Medium (MEM, Gibco 11095-080) with 10% FBS (Gibco 26140079), 1mM sodium pyruvate, and 0.1 mM non-essential amino acids in 5% CO_2_ and ambient O_2_ in a Forma series II CO_2_ incubator (Thermo). Fibroblast lines were transfected with an RNA-based reprogramming vector ReproRNA-OKSGM (Stemcell Technologies) containing *POU5F1*, *KLF4*, *SOX2*, *GLIS1*, and *c-MYC* according to the manufacturer’s protocol and incubated in 5% O_2_, 5% CO_2_, and ambient atmospheric pressure in an AVATAR cell control system (Xcell Biosciences). Single colonies were picked to establish iPSC lines and maintained in either 5% or 15% O_2_ and expanded for 3 passages before initiation of the experiment outlined in Figure 2A. iPSC-1 and iPSC-2 were generated from fibroblast lines GM04504 and GM00730, respectively. iPSC-3 was obtained from the NIH under the identifier NCRM-5 (male, cord-blood source).

### iPSC culture

All iPSC lines were maintained in mTesr1 (Stemcell Technologies) on ESC-grade reduced growth factor basement membrane matrix (Geltrex, Gibco) in the following incubator conditions using the AVATAR cell control system (Xcell Biosciences): 5% O_2_ + 2 PSI, 5% O_2_ + 0 PSI, 15% O_2_ + 2 PSI, and 15% O_2_ + 0 PSI all at 37° C and 5% CO_2_. iPSCs lines were passaged every 5 days using Gentle Cell Dissociation Reagent (Stemcell Technologies) and plated in mTesr1 in the presence of 10 uM Y-27632 (Stemcell Technologies). Medium was exchanged daily.

### NSC culture

Neural stem cell lines from cord-blood derived iPSCs (XCL-6, Xcell Science) and bone marrow derived iPSCs (ACS-5003, ATCC) were cultured on ESC-grade reduced growth factor basement membrane matrix (Geltrex, Gibco) in NSC maintenance medium (Xcell Science) and medium exchanged every other day.

### Differentiation to neural-ectoderm

For neural-ectoderm induction, iPSCs were washed 1x in PBS and dissociated into single-cell suspensions using Gentle Cell Dissociation Reagent (Stemcell Technologies) for 10 min at 37° C and seeded on to Geltrex-coated 12-well plates at a density of 700,000 cells in neural induction medium + 10 uM Y-27632 (Stemcell Technologies) and cultured for 7 days with daily medium exchange. 10 uM Y-27632 was omitted from the neural induction medium after day 2.

Neural induction medium (adapted from Li et al, 2011) is composed of 1:1 ratio of Advanced DMEM/F12 (Gibco) and Neurobasal (Gibco), 1x B27 (Gibco), 1x N2 (Gibco), 1x Glutamax (Gibco), 10 ng/mL human LIF (Peprotech), and 10 ng/mL human NOG (Peprotech).

### CNS-type neuronal differentiation

NSC lines were dissociated into single-cell suspensions using Accutase (Stemcell Technologies) and plated onto Poly-L-Ornithine (15 ug/mL, Sigma) / Laminin (10 ug/mL, Sigma) coated 12-well plates. NSC suspensions were seeded at 200,000 cells per well in STEMdiff Neuron Differentiation Kit medium (Stemcell Technologies) and medium replaced every other day until day 7. To initiate neuronal maturation, cells were dissociated into single-cell suspension using Accutase and re-seeded onto fresh Poly-L-Ornithine / Laminin coated 12-well plates at a density of 150,000 cells in STEMdiff Neuron Maturation Kit medium (Stemcell Technologies) and medium exchanged every other day until days 14 and 21.

### Motor neuron differentiation

We used the protocol described by Maury et al, 2015 with slight modifications. Briefly, 10,000 NSC cells (XCL-6 and ACS-5003) were seeded in each well of 96-well round-bottom suspension plates (Corning) in motor neuron differentiation medium with 3 uM CHIR99021 (Stemcell Technologies) and cultured for 2 days to form neurospheres. Medium was replaced with motor neuron differentiation medium containing 3 uM CHIR99021, 500 nM SAG (Tocris), and 100 nM retinoic acid (Sigma) and cultured for a further 2 days. Medium was then replaced with motor neuron differentiation medium containing only 500 nM SAG and 100 nM retinoic acid and cultured until day 9, with medium exchanged every other day. On day 9, neurospheres were pooled and treated with Accutase for 5-10 min at 37° C and gently pipetted to generate a single-cell suspension. 150-200,000 cells were seeded onto Poly-L-Ornithine (20 ug/mL) / Laminin (5 ug/mL) coated 12-well plates in motor neuron differentiation medium containing 10 uM DAPT and cultured for 2 days. On days 11, medium was replaced with motor neuron maturation medium and replaced every other day until day 31. Motor neuron differentiation medium is composed of 1:1 ratio of DMEM/F12 (Gibco) and Neurobasal (Gibco), 2% B27 (Gibco), 1% N2 (Gibco), 1x non-essential amino acids (Gibco), 1x Glutamax (Gibco), and 0.5 uM ascorbic acid (Sigma). Motor neuron maturation medium contains the same formulation as differentiation medium but with the addition of 1 uM db-cAMP (Sigma), 20 ng/mL human BDNF and 10 ng/mL human GDNF (both from Peprotech).

### HSPC differentiation

HSPC differentiation protocol and medium was adapted from Angelos et al, 2017. Briefly, fibroblast and cord blood derived iPSC lines (Line 7, GM00730 and NCRM-5, respectively) were seeded into 12-well plates coated with Geltrex at a concentration of 100 aggregates per well of approximately 100-200 nm in diameter in mTesr1 + 10 uM Y-27632 and incubated overnight in a CO_2_ incubator. On day 1, a full medium exchange with 1 mL of phase 1 medium containing 20 ng/mL human BMP4, 20 ng/mL human VEGF_121_, 40 ng/mL human SCF, 1% anti-anti (Gibco), and 5% PFHM-II (Gibco) in APEL 2 base medium (Stemcell Technologies). On day 3, a full medium exchange with 1 mL of fresh phase 1 medium was performed. On day 6, a full medium exchange was performed with phase 2 medium containing 40 ng/mL human VEGF_121_, 40 ng/mL human SCF, 30 ng/mL human TPO, 30 ng/mL human IL-3, 30 ng/mL human IL-6, 1% anti-anti, and 5% PFHM-II in APEL 2 base medium. On days 9 and 12, 80% of medium was removed followed by addition of 1.5 mL of fresh phase 2 medium. On day 14, supernatant was removed and stained for CD34, CD43, CD45, and Propidium Iodide (Life Technologies). Primary antibodies are listed in the table below. All proteins were purchased from Peprotech.

### Whole transcriptome mRNA-seq

Total RNA was purified using the Single-cell RNA Purification Kit (Norgen Biotek) and mRNA-seq libraries were prepared using TruSeq Stranded mRNA HS protocol (Illumina) using 500 ng total RNA input. Library quality was confirmed using the High Sensitivity DNA chip on the Agilent 2100 Bioanalyzer System (Agilent Technologies). mRNA-seq libraries were then run using 75 bp paired-end reads on a NextSeq 500 (Illumina).

### mRNA-seq analysis

Reads were aligned using hisat2 (PMID=25751142) v2.0.4 to the GRCh38 snp_trans reference genome downloaded from the hisat2 website. The resulting BAM file was processed using CleanSam from the picard v2.0.1 package (http://broadinstitute.github.io/picard) and sorted using samtools v1.3 (PMID:19505943). Read and alignment quality was evaluated using FastQC (http://www.bioinformatics.babraham.ac.uk/projects/fastqc/) and RSeQC v2.6.3 (PMID:22743226). Gene read counts were quantified using featureCounts v1.5.0-p1 (PMID:24227677) in paired-end mode and gene expression quantified using DESeq2 v1.14.0. Differential expression analysis was performed using DESeq2 v1.14.0.

For hierarchical clustering of mRNA-seq expression values and Spearman correlation values we used Cluster 3.0: [ref: https://www.ncbi.nlm.nih.gov/pubmed/14871861?dopt=AbstractPlus] with average linkage un-centered hierarchical clustering followed by heatmap visualization using Java TreeView v1.1.6r4 [ref: https://academic.oup.com/bioinformatics/article/20/17/3246/186177]. For figure 2B, a filtration step was applied to remove oxygen-specific significant differentially expressed genes extracted from DEseq2 analysis of 15% O_2_ + 0 PSI *vs*. 5% O_2_ + 0 PSI expression counts data of passage seven iPSCs.

GO term enrichment was performed using the goseq v3.8 package found here: (https://bioconductor.org/packages/release/bioc/html/goseq.html).

### Immunofluorescence microscopy

iPSC lines were plated onto glutaraldehyde-activated glass coverslips coated with ESC-grade reduced growth factor basement membrane matrix (Geltrex, Gibco). Neuronal cultures were plated onto glutaraldehyde-activated glass coverslips coated with appropriate substrates as mentioned above. Cells were fixed in 4% formaldehyde for 10 min and blocked in a solution containing 5% normal goat or donkey serum and 0.3% Triton X-100 in PBS without Ca^2+^/Mg^2+^. Primary antibodies were stained overnight at 4° C in antibody dilution buffer containing 1% BSA and 0.3% Triton X-100 in PBS without Ca^2+^/Mg^2+^. Primary antibody was washed 2x in PBS without Ca^2+^/Mg^2+^ and stained with DAPI and Alexa-conjugated IgG secondary antibodies for 1 hr at RT. Secondary antibody was washed 3x in PBS without Ca^2+^/Mg^2+^ and mounted with VECTASHIELD Hardset Antifade Mounting Medium (Vector #H-1400). Images were acquired on a fluorescence microscope (Zeiss) and all analysis performed using ImageJ with a minimum of 3 images per stain for each replicate experiment.

### Flow cytometry

For staining of neural-ectoderm cells for PAX6 and NES, cells were first stained with Live/Dead Fixable Violet Dead Cell Stain Kit (Invitrogen) then fixed in 2% formaldehyde for 10 min and washed 2x in PBS without Ca^2+^/Mg^2+^. Cells were permeabilized in 0.7% Tween-20 (Fisher) in PBS for 15 min at RT and washed 1x in PBS. Primary antibody staining was performed in a solution of 0.5% Tween-20, 1% BSA, and 10% normal goat serum in PBS for 1 hr at RT. Secondary antibody staining was performed at 1:1000 in PBS for 30 min in the dark, then washed 2x in PBS.

For CD34, CD43, CD45, and CD57 extra-cellular staining, conjugated primary antibodies were diluted in PBS without Ca^2+^/Mg^2+^ with 1% BSA and incubated with cells for 1 hr at 4° C, washed 2x in PBS, then analyzed on LSRII flow cytometer (BD Bioscience). Data analysis was performed using FlowJo v10.5.

### Primary antibodies

**Primary antibodies:**
| **Antibody name** | **Manufacturer / Catalog** | **Dilution** |
| --- | --- | --- |
| POU5F1 (Oct-4a) | Cell Signaling / 2840 | 1:400 |
| NANOG | Thermo Fisher / MA1-017 | 1:100 |
| SOX2 | Abcam / ab97959 | 1:400 |
| PAX6 | Biolegend / 901301 | 1:300 |
| NES (Nestin) | Thermo Fisher / MA1-110 | 1:100 |
| ChAT | Millipore / AB144P | 1:200 |
| SMI-32<br>(Neurofilament-H) | Biolegend / 801701 | 1:1000 |
| MAP2 | Aves Labs / MAP | 1:500 |
| SYN1 | Invitrogen / A6442 | 1:500 |
| CD57 | Sigma / C6680 | 1:30 |
| CD34 | BD / 562577 | 1:20 |
| CD43 | Stemcell Technologies / 60085PE | 1:20 |
| CD45 | Stemcell Technologies / 60018AD | 1:20 |

### Human fibroblast lines

**Human fibroblast lines:**
| **Cat #** | **Vendor** | **Gender** | **Age** |
| --- | --- | --- | --- |
| GM23962 | Coriell | Male | 27 YR |
| GM23963 | Coriell | Male | 48 YR |
| GM04504 | Coriell | Female | 32 YR |
| GM06111 | Coriell | Female | 16 FW |
| GM06168 | Coriell | Male | 20 FW |
| GM05294 | Coriell | Male | 22 YR |
| GM00730 | Coriell | Female | 45 YR |
| GM08399 | Coriell | Female | 19 YR |
| CRL-2097 | ATCC | Male | Neo-natal |
| CRL-2129 | ATCC | Female | 34 YR |

\* FW = fetal weeks

## Author Contributions

ZP and BA conceived the study. ZP designed and performed the experiments, wrote the manuscript. JL and BA reviewed and helped prepare the manuscript. ZP and BD performed the bioinformatics of the sequencing data.

## Acknowledgements

This work was funded by the California Institute of Regenerative Medicine (CIRM EDUC2-08391 to ZP)

## References

Angelos MG, Ruh PN, Webber BR, Blum RH, Ryan CD, Bendzick L, Shim S, Yingst AM, Tufa DM, Verneris MR, Kaufman DS. 2017. Aryl hydrocarbon receptor inhibition promotes hematolymphoid development from human pluripotent stem cells. Blood. 129(26):3428–3439.

Baba A, Ohtake F, Okuno Y, Yokota K, Okada M, Imai Y, Ni M, Meyer CA, Igarashi K, Kanno J, Brown M, Kato S. 2011. PKA-dependent regulation of the histone lysine demethylase complex PHF2–ARID5B. Nat Cell Biol. 13(6):668–75.

Battista S, Guarnieri D, Borselli C, Zeppetelli S, Borzacchiello A, Mayol L, Gerbasio D, Keene DR, Ambrosio L, Netti PA. 2005. The effect of matrix composition of 3D constructs on embryonic stem cell differentiation, Biomaterials. 26(31): 6194–6207.

Boroviak T, Loos R, Lombard P, Okahara J, Behr R, Sasaki E, Nichols J, Smith A, and Bertone P. 2015. Lineage-Specific Profiling Delineates the Emergence and Progression of Naive Pluripotency in Mammalian Embryogenesis. Developmental Cell. 35(3), 366–82.

Boulter E, Estrach S, Tissot FS, Hennrich ML, Tosello L, Cailleteau L, de la Ballina LR, Pisano S, Gavin AC, and Féral CC. 2018. Cell metabolism regulates integrin mechanosensing via an SLC3A2-dependent sphingolipid biosynthesis pathway. Nature Communications. 9(1), 4862.

Chan YS, Göke J, Ng JH, Lu X, Gonzales KA, Tan CP, Tng WQ, Hong ZZ, Lim YS, and Ng HH. 2013. Induction of a Human Pluripotent State with Distinct Regulatory Circuitry that Resembles Preimplantation Epiblast. Cell Stem Cell. 13(6): 663–675.

Chanson L, Brownfield D, Garbe JC, Kuhn I, Stampfer MR, Bissell MJ and LaBarge MA. 2011. Self-organization is a dynamic and lineage-intrinsic property of mammary epithelial cells. Proc. Natl. Acad. Sci. USA. 108, 3264–3269.

Chistiakov DA, Orekhov, AN, and Bobryshev YV. 2017. Effects of shear stress on endothelial cells: go with the flow. Acta Physiol. 219: 382–408.

Chow DC, Wenning LA, Miller WM, Papoutsakis TE. 2001. Modeling pO2 Distributions in the Bone Marrow Hematopoietic Compartment. II. Modified Kroghian Models. Biophysical Journal. 81(2): 685–696.

Collier AJ, Panula SP, Schell JP, Chovanec P, Plaza Reyes A, Petropoulos S, Corcoran AE, Walker R, Douagi I, Lanner F, and Rugg-Gunn PJ. 2017. Comprehensive Cell Surface Protein Profiling Identifies Specific Markers of Human Naive and Primed Pluripotent States. Cell Stem Cell. 20(6), 874–890.e7.

Correia C, Serra M, Espinha N, Sousa M, Brito C, Burkert K, Zheng Y, Hescheler J, Carrondo M, Sarić T, and Alves P. 2014. Combining Hypoxia and Bioreactor Hydrodynamics Boosts Induced Pluripotent Stem Cell Differentiation Towards Cardiomyocytes. Stem Cell Rev and Rep. 10, 786–801.

Covello KL, Kehler J, Yu H, Gordan JD, Arsham AM, Hu CJ, Labosky PA, Simon MC, and Keith B. 2006. HIF-2alpha regulates Oct-4: effects of hypoxia on stem cell function, embryonic development, and tumor growth. Genes and Development. 20(5), 557–70.

Eiraku M, and Sasai Y. 2012. Self-formation of layered neural structures in three-dimensional culture of ES cells. Curr. Opin. Neurobiol. 22, 768–777.

Engler A, Sen S, Sweeney H, and Discher D. 2006. Matrix Elasticity Directs Stem Cell Lineage Specification. Cell. 126(4), 677–689.

Formigli L, Meacci E, Sassoli C, Squecco R, Nosi D, Chellini F, Naro F, Francini F, and Zecchi-Orlandini S. 2007. Cytoskeleton/stretch-activated ion channel interaction regulates myogenic differentiation of skeletal myoblasts. J. Cell. Physiol., 211: 296–306.

Gao X, Tate P, Hu P, Tjian R, Skarnes WC, Wang Z. 2008. ES cell pluripotency and germ-layer formation require the SWI/SNF chromatin remodeling component BAF250a. Proceedings of the National Academy of Sciences. 105 (18): 6656–6661.

Giese AK, Frahm J, Hübner R, Luo J, Wree A, Frech MJ, Rolfs A, and Ortinau A. 2010. Erythropoietin and the effect of oxygen during proliferation and differentiation of human neural progenitor cells. BMC Cell Biol. 11, 94.

Hawkins RD, Hon GC, Lee LK, Ngo Q, Lister R, Pelizzola M, Edsall LE, Kuan S, Luu Y, Klugman S, Antosiewicz-Bourget J, Ye Z, Espinoza C, Agarwahl S, Shen L, Ruotti V, Wang W, Stewart R, Thomson JA, Ecker JR, and Ren B. 2010. Distinct epigenomic landscapes of pluripotent and lineage-committed human cells. Cell Stem Cell. 6(5):479–91.

Hong S-H, Werbowetski-Ogilvie T, Ramos-Mejia V, Lee JB, and Bhatia M. 2010. Multiparameter comparisons of embryoid body differentiation toward human stem cell applications. Stem Cell Research. 5(2):120–130.

Ivanovic Z, Hermitte F, Brunet de la Grange P, Dazey B, Belloc F, Lacombe F, Vezon G, and Praloran Gl. 2004. Simultaneous maintenance of human cord blood SCID-repopulating cells and expansion of committed progenitors at low O_2_ concentration (3%). Stem Cells. 2004. 22:716–24.

Kang J, Lienhard M, Pastor WA, Chawla A, Novotny M, Tsagaratou A, Lasken RS, Thompson EC, Surani MA, Koralov SB, Kalantry S, Chavez L, and Rao A. 2015. Simultaneous deletion of the methylcytosine oxidases Tet1 and Tet3 increases transcriptome variability in early embryogenesis. Proc Natl Acad Sci. 112(31), E4236–45.

Kao Y-C, Jheng J-R, Pan H-J, Liao W-Y, Lee C-H, and P-L Kuo. 2017. Elevated hydrostatic pressure enhances the motility and enlarges the size of the lung cancer cells through aquaporin upregulation mediated by caveolin-1 and ERK1/2 signaling. Oncogene. 36,863–874.

Kim JE, Lee EJ, Yanru Wu, Kang YG, and Shin J-W. 2019. The combined effects of hierarchical scaffolds and mechanical stimuli on ex vivo expansion of haematopoietic stem/progenitor cells. Artif Cell Nanomed Biotechnol. 47(1):585–592.

Kennedy M, Awong G, Sturgeon CM, Ditadi A, LaMotte-Mohs R, Zúñiga-Pflücker JC, and Keller G. 2012. T Lymphocyte Potential Marks the Emergence of Definitive Hematopoietic Progenitors in Human Pluripotent Stem Cell Differentiation Cultures. Cell Reports. 2(6):1722–1735.

Komatsu K, and Fujimori T. 2015. Multiple phases in regulation of Nanog expression during pre-implantation development. Develop. Growth Differ. 57: 648–656.

Labedz-Maslowska A, Szkaradek A, Mierzwinski T, Madeja Z, and Zuba-Surma E. 2021. Processing and Ex Vivo Expansion of Adipose Tissue-Derived Mesenchymal Stem/Stromal Cells for the Development of an Advanced Therapy Medicinal Product for use in Humans. Cells. 10, 1908.

Lee E, Kim J, Kang Y, and Shin JW. 2020. A Platform for Studying of the Three-Dimensional Migration of Hematopoietic Stem/Progenitor Cells. Tissue Eng. Regen. Med. 17(1):25–31.

Li H, Luo Q, Shan W, Cai S, Tie R, Xu Y, Lin Y, Qian P, and Huang H. 2021. Biomechanical cues as master regulators of hematopoietic stem cell fate. Cell Mol Life Sci. 78(16): 5881–5902.

Li W, Sun W, Zhang Y, Wei W, Ambasudhan R, Xia P, Talantova M, Lin T, Kim J, Wang X, Kim WR, Lipton SA, Zhang K, and Ding S. 2011. Rapid induction and long-term self-renewal of primitive neural precursors from human embryonic stem cells by small molecule inhibitors. Proc Natl Acad Sci 108(20), 8299–304.

Leipzig N, and Shoichet M. 2009. The effect of substrate stiffness on adult neural stem cell behavior. Biomaterials. 30; 6867–6878.

Lopes M, Belo I, and Mota M. 2014. Over-pressurized bioreactors: Application to microbial cell cultures. Biotechnology Progress. 30: 767–775.

Luo N, Conwell MD, Chen X, Kettenhofen CI, Westlake CJ, Cantor LB, Wells CD, Weinreb RN, Corson TW, Spandau DF, Joos KM, Iomini C, Obukhov AG, and Sun Y. 2014. Primary cilia signaling mediates intraocular pressure sensation. Proc Natl Acad Sci. 111(35):12871–6.

Mathieu J, Zhou W, Xing Y, Sperber H, Ferreccio A, Agoston Z, Kuppusamy K T, Moon RT, and Ruohola-Baker H. 2014. Hypoxia-inducible factors have distinct and stage-specific roles during reprogramming of human cells to pluripotency. Cell Stem Cell. 14(5), 592–605.

Mathieu J, Zhang Z, Nelson A, Lamba D, Reh T, Ware C, and Ruohola–Baker H. 2013. Hypoxia Induces Re-entry of Committed Cells into Pluripotency. Stem Cells. 31, 1737–1748.

Maury Y, Côme J, Piskorowski RA, Salah-Mohellibi N, Chevaleyre V, Peschanski M, Martinat C, and Nedelec S. 2015. Combinatorial analysis of developmental cues efficiently converts human pluripotent stem cells into multiple neuronal subtypes. Nature Biotechnology. 33, 89–96.

McKee C, and Chaudhry RG. 2017. Advances and challenges in stem cell culture. Colloids and Surfaces B: Biointerfaces. 159: 62–77.

Mulas C, Kalkan T, and Smith A. 2016. Nodal secures pluripotency upon embryonic stem cell progression from the ground state. Stem Cell Reports. 9(1): 77–91.

Murrow LM, Weber RJ, and Gartner ZJ. 2017 Dissecting the stem cell niche with organoid models: an engineering-based approach. Development. 144(6):998–1007.

Nichols J, and Smith A. 2009. Naive and primed pluripotent states. Cell Stem Cell. 4, 487–492.

Northey JJ, Przybyla L, and Weaver VM. 2017. Tissue Force Programs Cell Fate and Tumor Aggression. Cancer Discov. 7(11):1224–1237.

Ohashi T, Sugaya Y, Sakamoto N, and Sato M. 2007. Hydrostatic pressure influences morphology and expression of VE-cadherin of vascular endothelial cells. Journal of Biomechanics. 40(11): 2399–2405.

Ortega JA, Sirois CL, Memi F, Glidden N, and Zecevic N. 2016. Oxygen Levels Regulate the Development of Human Cortical Radial Glia Cells. Cereb Cortex. 27(7):3736–3751.

Prigione A, and Adjaye J. 2010. Modulation of mitochondrial biogenesis and bioenergetic metabolism upon in vitro and in vivo differentiation of human ES and iPS cells. Int. J. Dev. Biol. 54: 1729–1741.

Raghavan S, and Bitar KN. 2014. The influence of extracellular matrix composition on the differentiation of neuronal subtypes in tissue engineered innervated intestinal smooth muscle sheets. Biomaterials. 35(26):7429–40.

Runswick SK, O’Hare MJ, Jones L, Streuli CH, and Garrod DR. 2001. Desmosomal adhesion regulates epithelial morphogenesis and cell positioning. Nat. Cell Biol. 3, 823–830.

Saha K, Keung AJ, Irwin EF, Li Y, Little L, Schaffer DV, and Healy KE. 2008. Substrate modulus directs neural stem cell behavior. Biophysical journal, 95(9), 4426–38.

Sakadzić S, Roussakis E, Yaseen MA, Mandeville ET, Srinivasan VJ, Arai K, Ruvinskaya S, Devor A, Lo EH, Vinogradov SA, and Boas DA. 2010. Two-photon high-resolution measurement of partial pressure of oxygen in cerebral vasculature and tissue. Nat Methods. 7(9):755–759.

Shimizu N, Yamamoto K, Obi S, Kumagaya S, Masumura T, Shimano Y, Naruse K, Yamashita JK, Igarashi T, and Ando J. 2008. Cyclic strain induces mouse embryonic stem cell differentiation into vascular smooth muscle cells by activating PDGF receptor beta. J Appl Physiol. 104:766–72.

Spencer JA, Ferraro F, Roussakis E, Klein A, Wu J, Runnels JM, Zaher W, Mortensen LJ, Alt C, Turcotte R, Yusuf R, Cote D, Vinogradov SA, Scadden DT, and Lin CP. 2014. Direct measurement of local oxygen concentration in the bone marrow of live animals. Nature. 508(7495):269–273.

Stanley AC, Lounsbury KM, Corrow K, Callas PW, Zhar R, Howe AK, and Ricci MA. 2005. Pressure elevation slows the fibroblast response to wound healing. Journal of Vascular Surgery. 42(3): 546–551.

Studer L, Csete M, and Lee SH. 2000 Enhanced Proliferation, Survival, and Dopaminergic Differentiation of CNS Precursors in Lowered Oxygen. Neuroscience. 20, 7377–7383.

Timmermans F, Velghe I, Vanwalleghem L, Smedt MD, Coppernolle SV, Taghon T, Moore HD, Leclercq G, Langerak AW, Kerre T, Plum J, and Vandekerckhove B. 2009. Generation of T Cells from Human Embryonic Stem Cell-Derived Hematopoietic Zones. The Journal of Immunology. 182(11): 6879–6888.

Tworkoski E, Glucksberg MR, and Johnson M. 2018. The effect of the rate of hydrostatic pressure depressurization on cells in culture. PLoS ONE. 13 (1).

Varum S, Rodrigues AS, Moura MB, Momcilovic O, Easley CA, Ramalho-Santos J, Van Houten B, and Schatten G. 2011. Energy metabolism in human pluripotent stem cells and their differentiated counterparts. PloS one, 6(6), e20914.

Wagner DR, Lindsey DP, Li KW, Tummala P, Chandran SE, Smith RL, Longaker AT, Carter DR, and Beaupre JS. 2008. Hydrostatic pressure enhances chondrogenic differentiation of human bone marrow stromal cells in osteochondrogenic medium. Ann Biomed Eng. 36, 813–820.

Wang X, Zhang Z, Tao H, Liu J, Hopyan S, and Sun Y. 2018. Characterizing Inner Pressure and Stiffness of Trophoblast and Inner Cell Mass of Blastocysts. Biophysical Journal. 115, 2443–2450.

Wolfe RP, and Ahsan T. 2013. Shear stress during early embryonic stem cell differentiation promotes hematopoietic and endothelial phenotypes. Biotechnol Bioeng. 110(4):1231–1242.

Wu H and Sun YE. 2006. Epigenetic regulation of stem cell differentiation. Pediatric Research. 59: 21R–25R.

Xie Y, Zhang J, Lin Y, Gaeta X, Meng X, Wisidagama DR, Cinkornpumin J, Koehler CM, Malone CS, Teitell MA, and Lowry WE. 2014. Defining the role of oxygen tension in human neural progenitor fate. Stem Cell Reports. 3(5):743–57.

Yan L, Yang M, Guo H, Yang L, Wu J, Li R, Liu P, Lian Y, Zheng X, Yan J, Huang J, Li M, Wu X, Wen L, Lao K, Li R, Qiao J, and Tang F. 2013. Single-cell RNA-Seq profiling of human preimplantation embryos and embryonic stem cells. Nat. Structural Mol. Biol. 20: 1131–1139.

Yasui T, Uezono N, Nakashima H, Noguchi H, Matsuda T, Noda-Andoh T, Okano H, and Nakashima K. 2017. Hypoxia Epigenetically Confers Astrocytic Differentiation Potential on Human Pluripotent Cell-Derived Neural Precursor Cells. Stem Cell Reports. 8(6): 1743–1756.

Yin X, Mead BE, Safaee H, Langer R, Karp JM, and Levy O. 2016. Engineering Stem Cell Organoids. Cell Stem Cell. 18(1):25–38.

Yoshida Y, Takahashi K, Okita K, Ichisaka T, and Yamanaka S. 2009. Hypoxia Enhances the Generation of Induced Pluripotent Stem Cells. Cell Stem Cell. 5, 237–241.

Zhao Y-H, Lv X, Liu Y-L, Zhao Y, Li Q, Chen Y-J, Zhang M. 2015. Hydrostatic pressure promotes the proliferation and osteogenic/chondrogenic differentiation of mesenchymal stem cells: The roles of RhoA and Rac1. Stem Cell Research. 14(3): 283–296.

Zhong X, Gutierrez C, Xue T, Hampton C, Vergara MN, Cao LH, Peters A, Park TS, Zambidis ET, Meyer JS, Gamm DM, Yau KW, and Canto-Soler MV. 2014. Generation of three-dimensional retinal tissue with functional photoreceptors from human iPSCs. Nature Communications, 5, 4047.

Zhu L.L., Wu LY, Yew DT, and Fan M. 2005. Effects of Hypoxia on the Proliferation and Differentiation of NSCs. Mol. Neurobiol., 31, 231–242.

