## Supplementary material for "Application of Elevated Atmospheric Pressure and Hypoxia Enhance Pluripotency and Stem Cell Differentiation": Supplmental Files

#### Supplemental Figure Legends:

##### Figure S1.

Day 19 post-transfection re-programming results for CRL-2097 fibroblast line in the indicated oxygen and atmospheric pressure conditions. Data is presented as the fold-change in re-programming efficiency compared to 5% $O_2$ + 0 PSI. Re-programming efficiency is calculated as the number of iPSC colonies / number of transfected fibroblasts. n = 4 independent re-programmings except for 5% $O_2$ + 5 PSI, which was performed as n = 2. Error bars are standard error of the mean.

##### Figure S2.

**(a)** Flow cytometry analysis of passage three iPSCs cultured in 5% $O_2$ stained for the primed-state pluripotency associated surface marker CD57 (B3GAT1).
**(b)** Glycolysis and oxidative phosphorylation pathway-related genes from Hallmark, GO, and Kegg gene ontology databases that are differentially expressed between 5% $O_2$ + 2 PSI *vs*. 5% $O_2$ + 0 PSI at passage three with adjusted (FDR) p-value < 0.05. Un-supervised hierarchical clustering of genes / samples shown as scaled log2 fold change across rows (genes).
**(c)** Box-plot of cumulative significant differentially expressed genes from **(b)** above and **(4d)** with 5% $O_2$ + 2 PSI *vs*. 5% $O_2$ + 0 PSI genes included. The total number of significant genes is listed above each group. p-values determined by Wilcox Rank-sum test. * = p-value < 0.05, ** = < 0.01.
**(d)** The top 10 GO terms for down-regulated differentially expressed genes between 5% $O_2$ + 2 PSI *vs*. 5% $O_2$ + 0 PSI at passage three and seven. Gene ontology enrichment ranked according to FDR adjusted p-value and displayed on log10 scale. The number of significant differentially expressed genes for each GO term is displayed in the bar graph.

##### Figure S3.

**(a)** Principal component analysis of whole-transcriptome mRNA-seq expression data for iPSCs at passage three and seven in the indicated culture conditions.

**(b)** Representative immunofluorescence images of iPSCs at passage three showing expression of POU5F1 and NANOG in the indicated culture conditions. Scale bar = 50 µm.

**Figure S4.**

Heatmap showing overlapping genes at passage three that are significantly differentially expressed in response to elevated pressure in both 5% and 15% $O_2$ with an FDR adjusted p-value < 0.05 between 15% $O_2$ + 2 PSI *vs.* 15% $O_2$ + 0 PSI and 5% $O_2$ + 2 PSI *vs.* 5% $O_2$ + 0 PSI using DEseq2.

Figure S1

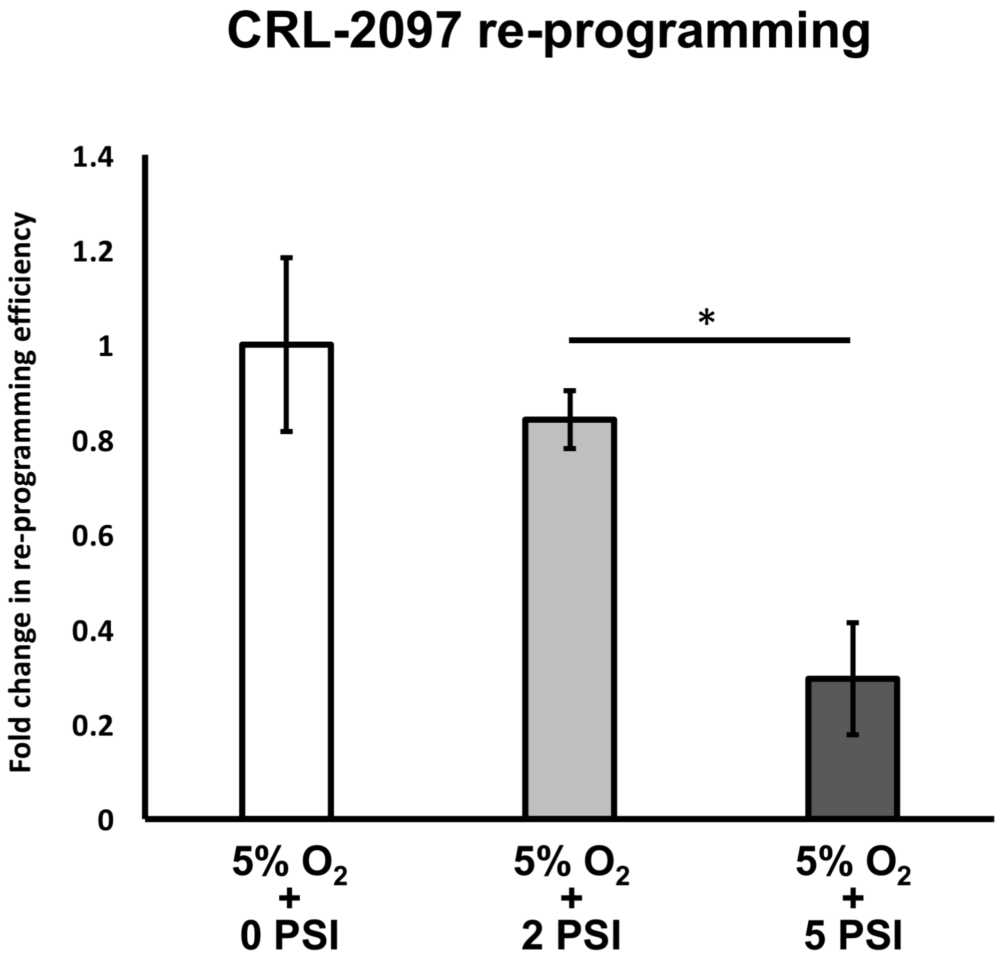
CRL-2097 re-programming
Fold change in re-programming efficiency
1.4
1.2
1
0.8
0.6
0.4
0.2
0
*
5% O2 + 0 PSI
5% O2 + 2 PSI
5% O2 + 5 PSI

### Figure S2

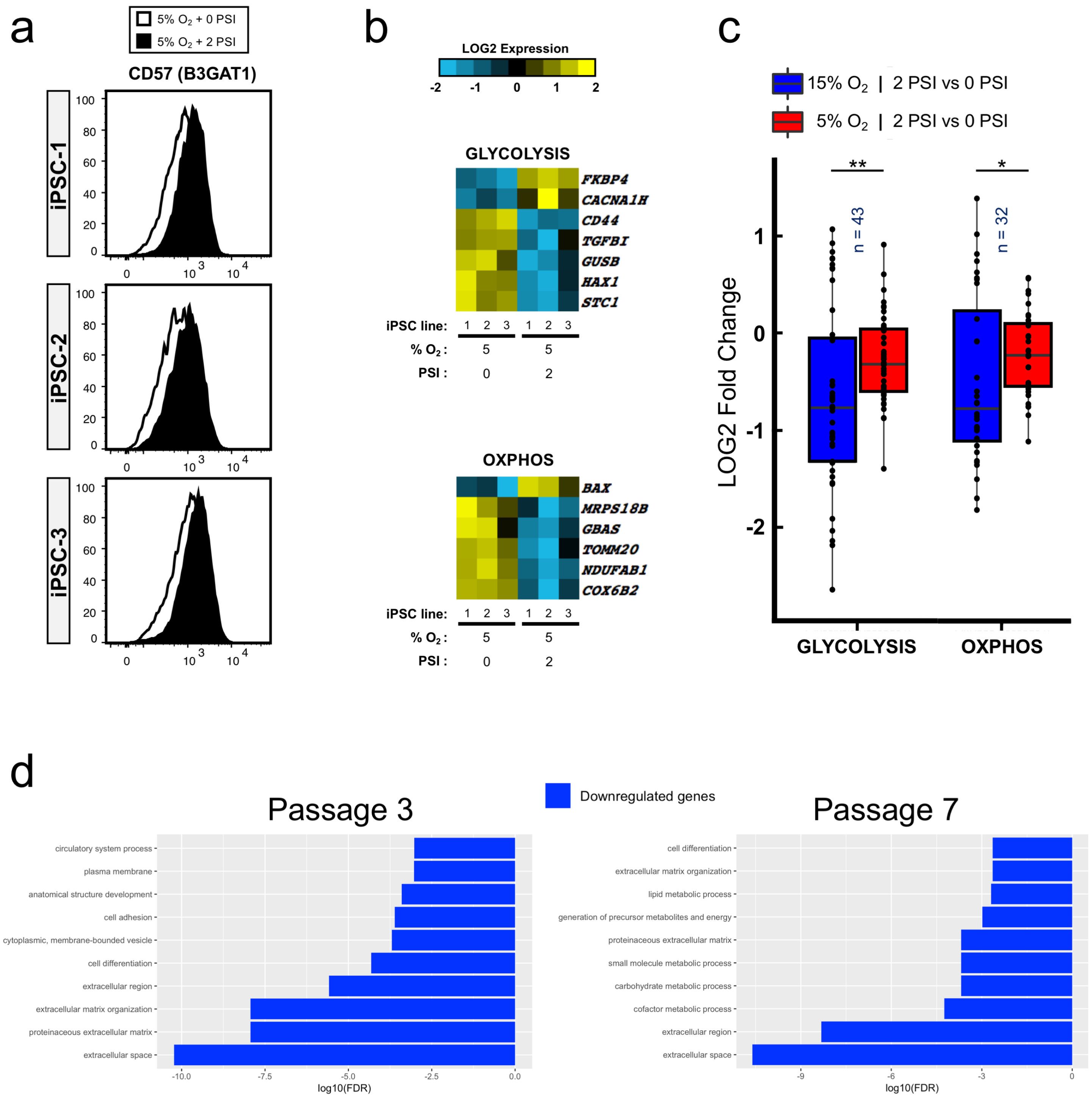

a
5% $O_2$ + 0 PSI
5% $O_2$ + 2 PSI
CD57 (B3GAT1)
iPSC-1
iPSC-2
iPSC-3
100
80
60
40
20
0
0
$10^3$
$10^4$
b
LOG2 Expression
-2
-1
0
1
2
GLYCOLYSIS
FKBP4
CACNA1H
CD44
TGFBI
GUSB
HAX1
STC1
iPSC line:
1 2 3 1 2 3
% $O_2$ :
5
5
PSI :
0
2
OXPHOS
BAX
MRPS18B
GBAS
TOMM20
NDUFAB1
COX6B2
c
15% $O_2$ | 2 PSI vs 0 PSI
5% $O_2$ | 2 PSI vs 0 PSI
LOG2 Fold Change
**
*
n = 43
n = 32
GLYCOLYSIS
OXPHOS
d
Passage 3
Downregulated genes
Passage 7
circulatory system process
plasma membrane
anatomical structure development
cell adhesion
cytoplasmic, membrane-bounded vesicle
cell differentiation
extracellular region
extracellular matrix organization
proteinaceous extracellular matrix
extracellular space
-10.0
-7.5
-5.0
-2.5
0.0
log10(FDR)
cell differentiation
extracellular matrix organization
lipid metabolic process
generation of precursor metabolites and energy
proteinaceous extracellular matrix
small molecule metabolic process
carbohydrate metabolic process
cofactor metabolic process
extracellular region
extracellular space
-9
-6
-3
0
log10(FDR)

### Figure S3

a

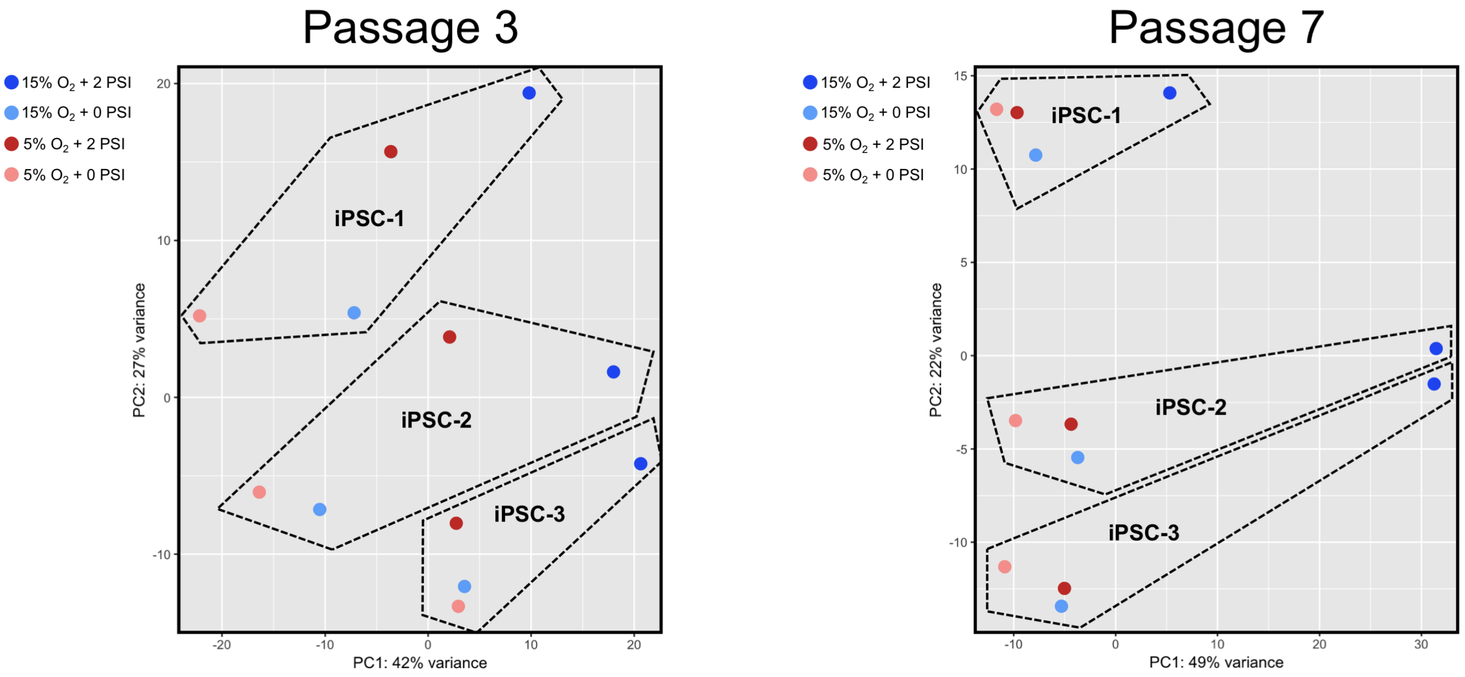

Passage 3
15% $O_2$ + 2 PSI
15% $O_2$ + 0 PSI
5% $O_2$ + 2 PSI
5% $O_2$ + 0 PSI
iPSC-1
iPSC-2
iPSC-3
PC1: 42% variance
PC2: 27% variance
Passage 7
PC1: 49% variance
PC2: 22% variance


b

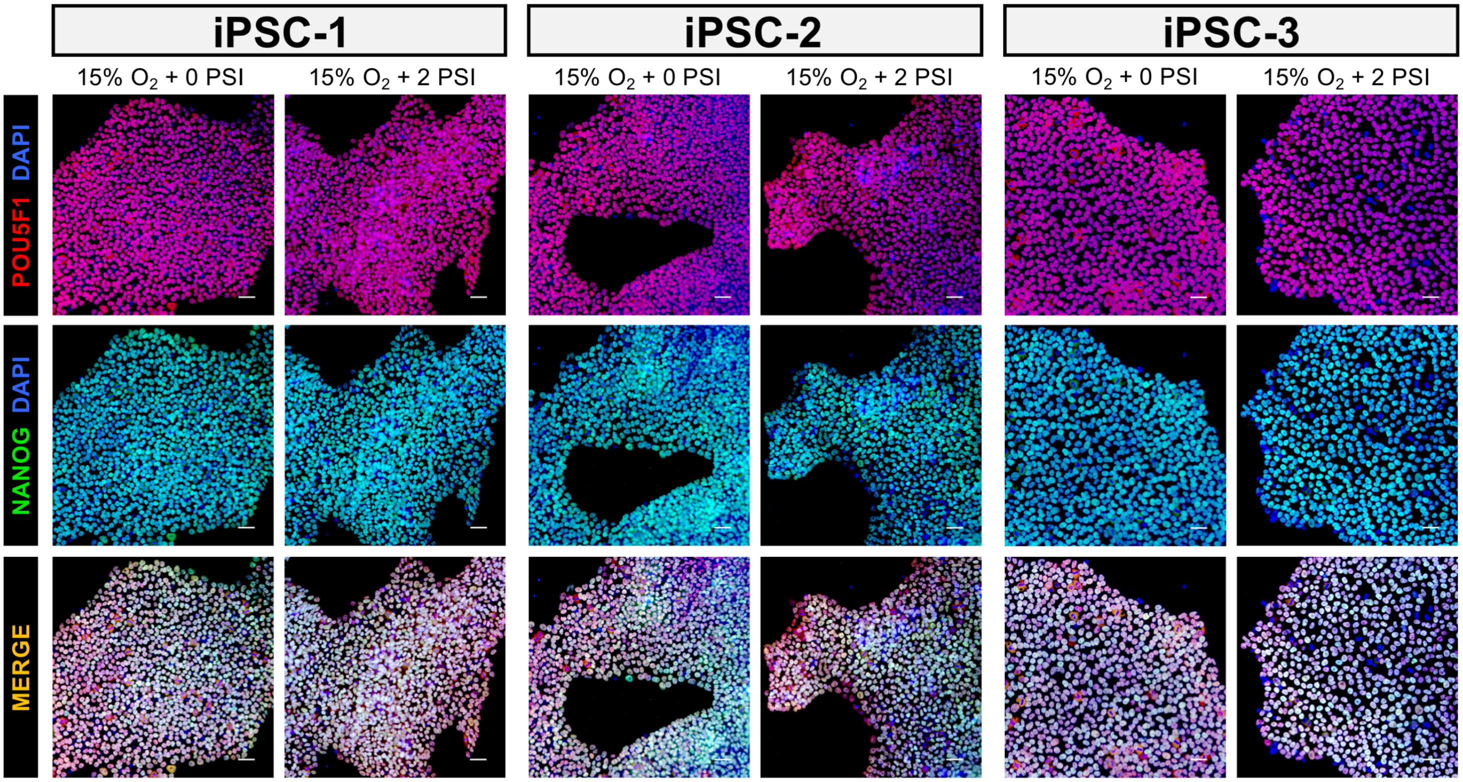

iPSC-1
iPSC-2
iPSC-3
15% $O_2$ + 0 PSI
15% $O_2$ + 2 PSI
POU5F1 DAPI
NANOG DAPI
MERGE

### Figure S4

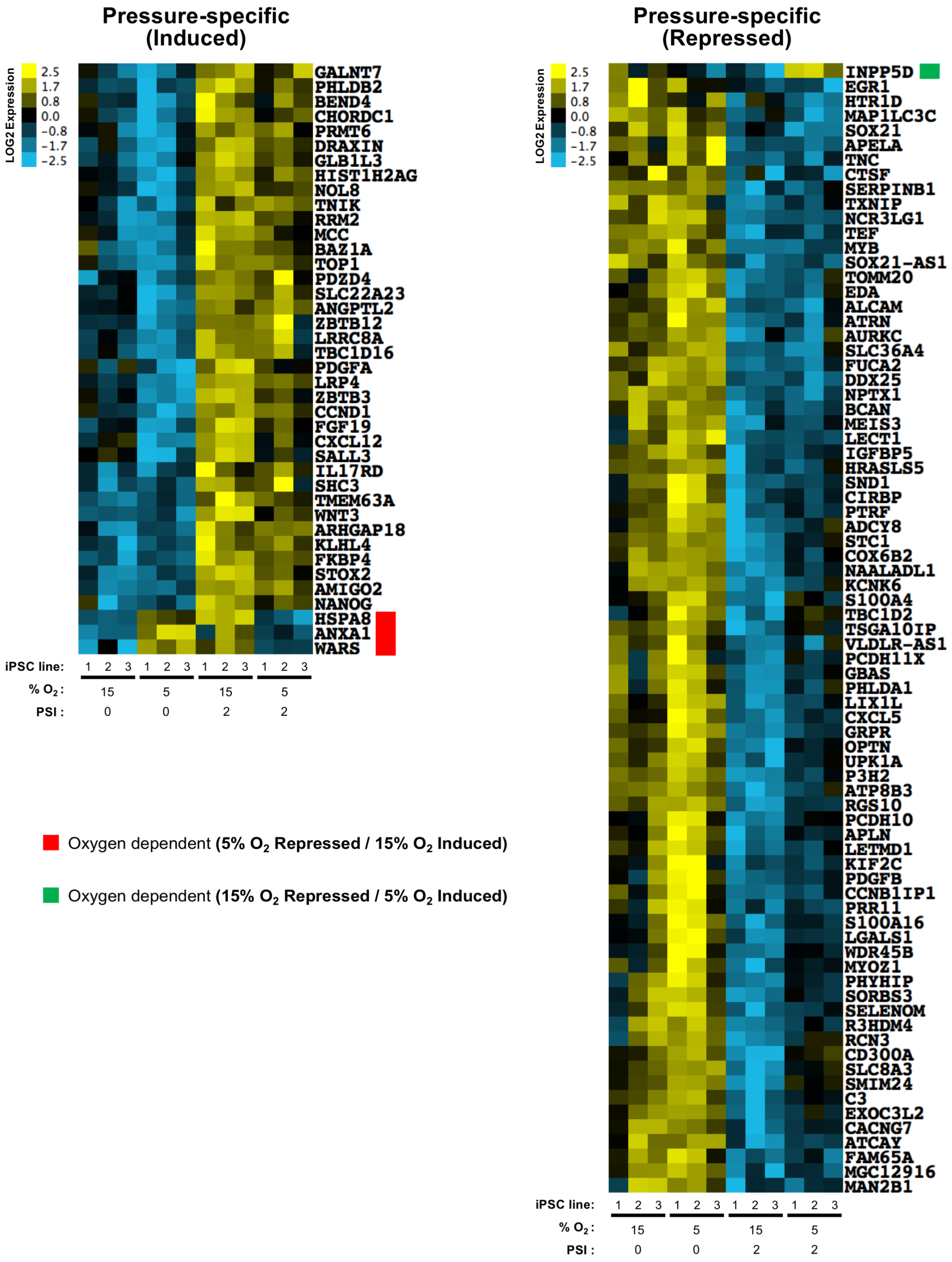
Pressure-specific (Induced)
LOG2 Expression
2.5
1.7
0.8
0.0
-0.8
-1.7
-2.5
GALNT7
PHLDB2
BEND4
CHORDC1
PRMT6
DRAXIN
GLB1L3
HIST1H2AG
NOL8
TNIK
RRM2
MCC
BAZ1A
TOP1
PDZD4
SLC22A23
ANGPTL2
ZBTB12
LRRC8A
TBC1D16
PDGFA
LRP4
ZBTB3
CCND1
FGF19
CXCL12
SALL3
IL17RD
SHC3
TMEM63A
WNT3
ARHGAP18
KLHL4
FKBP4
STOX2
AMIGO2
NANOG
HSPA8
ANXA1
WARS
iPSC line: 1 2 3 1 2 3 1 2 3 1 2 3
% $O_2$: 15 5 15 5
PSI : 0 0 2 2
Oxygen dependent (5% $O_2$ Repressed / 15% $O_2$ Induced)
Oxygen dependent (15% $O_2$ Repressed / 5% $O_2$ Induced)
Pressure-specific (Repressed)
LOG2 Expression
2.5
1.7
0.8
0.0
-0.8
-1.7
-2.5
INPP5D
EGR1
HTR1D
MAP1LC3C
SOX21
APELA
TNC
CTSF
SERPINB1
TXNIP
NCR3LG1
TEF
MYB
SOX21-AS1
TOMM20
EDA
ALCAM
ATRN
AURKC
SLC36A4
FUCA2
DDX25
NPTX1
BCAN
MEIS3
LECT1
IGFBP5
HRASLS5
SND1
CIRBP
PTRF
ADCY8
STC1
COX6B2
NAALADL1
KCNK6
S100A4
TBC1D2
TSGA10IP
VLDLR-AS1
PCDH11X
GBAS
PHLDA1
LIX1L
CXCL5
GRPR
OPTN
UPK1A
P3H2
ATP8B3
RGS10
PCDH10
APLN
LETMD1
KIF2C
PDGFB
CCNB1IP1
PRR11
S100A16
LGALS1
WDR45B
MYOZ1
PHYHIP
SORBS3
SELENOM
R3HDM4
RCN3
CD300A
SLC8A3
SMIM24
C3
EXOC3L2
CACNG7
ATCAY
FAM65A
MGC12916
MAN2B1
iPSC line: 1 2 3 1 2 3 1 2 3 1 2 3
% $O_2$: 15 5 15 5
PSI : 0 0 2 2
